# The estrous cycle modulates hippocampal spine dynamics, dendritic processing, and spatial coding

**DOI:** 10.1101/2024.08.02.606418

**Authors:** Nora S. Wolcott, William T. Redman, Marie Karpinska, Emily G. Jacobs, Michael J. Goard

## Abstract

Histological evidence suggests that the estrous cycle exerts a powerful effect on CA1 neurons in mammalian hippocampus. Decades have passed since this landmark observation, yet how the estrous cycle shapes dendritic spine dynamics and hippocampal spatial coding *in vivo* remains a mystery. Here, we used a custom hippocampal microperiscope and two-photon calcium imaging to track CA1 pyramidal neurons in female mice over multiple cycles. Estrous cycle stage had a potent effect on spine dynamics, with heightened density during periods of greater estradiol (proestrus). These morphological changes were accompanied by greater somatodendritic coupling and increased infiltration of back-propagating action potentials into the apical dendrite. Finally, tracking CA1 response properties during navigation revealed enhanced place field stability during proestrus, evident at the single-cell and population level. These results establish the estrous cycle as a driver of large-scale structural and functional plasticity in hippocampal circuits essential for learning and memory.

## Introduction

Circulating sex steroid hormones have a powerful, but poorly understood, impact on neuronal structure and function^1,2^. Receptors for two steroid hormones in particular, 17β-estradiol and progesterone, are highly expressed in the hippocampus^2,3^, an area critical for episodic and spatial memory formation^4–6^. These steroid hormones drive complex intracellular signaling cascades and, in humans, promote functional reorganization of brain networks across the reproductive cycle^1,7–9^. However, a circuit-level understanding of how hormones shape neuronal processing and plasticity is virtually unknown.

Foundational *ex vivo* studies in the early 1990s revealed that dendritic spines, the primary site of excitatory synapses, exhibit significant changes in density over the course of the 4-5 day estrous cycle in the rodent hippocampus, particularly in apical CA1 dendrites^10–14^. Later work showed that estradiol enhances excitatory synaptic plasticity by stimulating the insertion of NMDA receptors into the postsynaptic membrane of CA1 pyramidal neurons^15^. The synaptogenic effects of naturally circulating estradiol in CA1 have been observed in several mammalian species, including rats and non-human primates^10,12,16^. These effects are modulated by steroid hormone receptors that act through both genomic and non-genomic pathways, resulting in widespread transcriptional, translational, and epigenomic changes^17, 18^. Additionally, recent studies in humans found that rhythmic changes in steroid hormone production across the menstrual cycle are sufficient to drive widespread changes in functional connectivity across the cortical mantle, along with pronounced morphological changes in hippocampal subfields^7, 19, 20^. Together, this work cements sex steroid hormones as potent neuromodulators in the mammalian hippocampus, but we still lack a critical cellular and circuit-level understanding of how sex steroid hormones shape neural processing *in vivo*. Recent developments in multiphoton imaging techniques now allow us to address this knowledge gap using longitudinal observations of the same dendritic processes over unprecedented timescales^21^. This paves the way to answer long standing questions regarding the role of a cyclic endocrine environment on synaptic plasticity. For instance, are spines that emerge in response to heightened estradiol quickly pruned, or are they stable across multiple cycles? How does the addition or removal of spines influence how CA1 neurons integrate their excitatory inputs? Finally, how might these changes influence hippocampal responses and spatial coding?

Prior work offers indirect evidence that the estrous cycle may indeed have such effects. For example, new spines resulting from estradiol administration form functional synapses with new presynaptic partners^22^. This suggests that during high estradiol stages of the estrous cycle, CA1 neurons likely experience increased excitatory synaptic input from upstream neurons, which may exert a powerful influence on neuronal processing. As an example, these changes could shape the spatial coding of hippocampal place cells, a subset of CA1 neurons that encode specific locations in an animal’s environment^4, 23, 24^. The formation of place cells is thought to be highly dependent on local dendritic events^25^, so hormone-driven changes in excitability could reasonably influence the formation or stability of hippocampal place fields, with broad implications for spatial memory and navigational abilities.

Hippocampal place cells are highly adaptive; in response to a change in the spatial environment, such as visual cues, the shape of the chamber, or the reward location, the population of place cells rapidly remap their preferred spatial locations^26–29^. When reintroduced to the same environment, place cells regain their original firing location^26, 27^. These properties are highly evolutionarily conserved, and allow animals to successfully remember distinct environments^30–32^.

To address these questions, we used longitudinal two-photon imaging of naturally cycling female mice to determine the impact of the murine estrous cycle on CA1 neuron morphology and function. We found that dendritic spine dynamics, dendritic processing, and place cell stability all undergo pronounced estrous-dependent modulation. Taken together, these findings provide insight into how endocrine factors shape brain regions critical to spatial cognition, from the synapse to the circuit level.

## Results

### The estrous cycle modulates dendritic spine turnover and morphology

The rodent estrous cycle lasts 4-5 days and can be divided into the following stages: diestrus, proestrus, estrus, and metestrus^33–35^. The cycle follows a similar trend to that of the human 28- day menstrual cycle, in which estradiol levels rise through the follicular phase, corresponding to the diestrus and proestrus stages, before peaking to stimulate ovulation, after which estradiol levels fall precipitously at the beginning of the luteal phase, corresponding to the estrus stage (Fig. 1A)^7, 34^. The most common and reliable method for chronic, non-invasive estrous staging in rodents is vaginal cytology, in which epithelial cells are collected from the female mouse, and the relative proportion of cell types (typically cornified epithelial, nucleated epithelial, and leukocytes) is used to determine estrous stage^34, 35^. However, this method is subjective, and has been shown to suffer from inconsistencies between examiners^36, 37^. Here, estrous stage was confirmed by EstrousNet, a deep learning-based classifier of estrous stage from vaginal cytology, automating and streamlining the staging process (Fig. 1A)^36^. Classifications were performed after data collection and independently of image processing to blind the experimenter to estrous stage.

**Figure 1.**
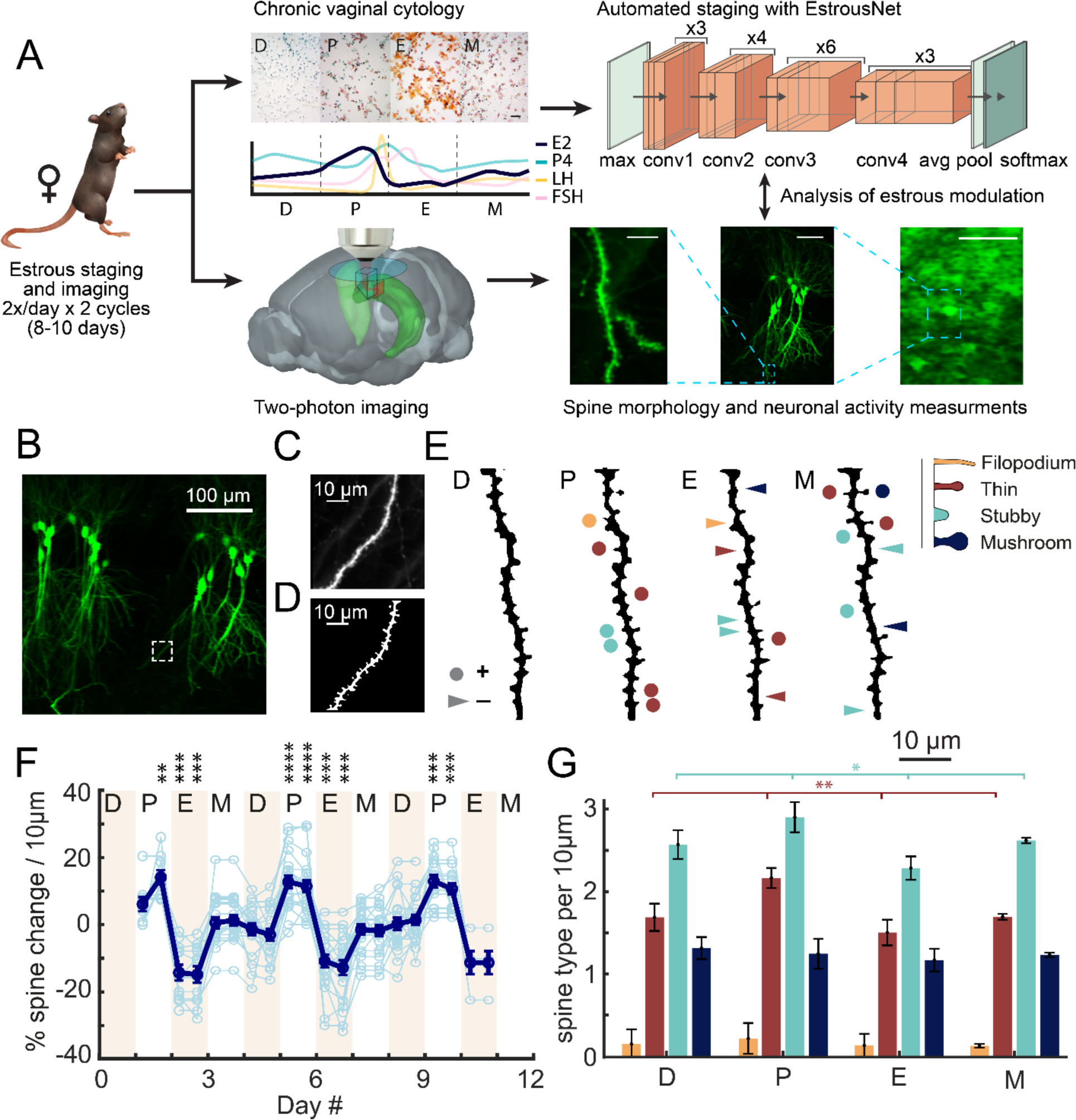
Dendritic spine density is longitudinally modulated by estrous cycle stage. A. Experimental pipeline schematic. Measurements were taken from female mice once every 12 hours across two consecutive estrous cycles (8-10 days). Prior to each session, vaginal lavage samples were taken and stained using Shorr stain (top). Estrous stage in upper left hand corner (D = diestrus, P = proestrus, E = estrus, M = metestrus). Scale bar = 1 mm. Each estrous stage is representative of a unique hormonal profile. Representation of the relative concentrations of ovarian hormones across the four estrous stages are shown, including 17β- estradiol (E2, navy), progesterone (P4, blue), luteinizing hormone (LH, yellow), and follicle-stimulating hormone (FSH, pink)^34^. Classifications were performed using EstrousNet, with transfer learning ResNet50 architecture shown here (top right), including four convolutional modules converging on a pooling and softmax classification layer. We then used two-photon imaging to track the structure or functional responses of hippocampal neurons (bottom) to determine changes across sessions. A schematic illustrating microperiscope implantation and light path for hippocampal imaging in a Thy1-GFP-M mouse demonstrates the imaging technique for dendritic experiments. Representative images are shown for dendritic spines (scale bar = 10 μm), whole dendrites (scale bar = 50 μm), and CA1 population imaging (scale bar = 100 μm). Finally, imaging data is matched with estrous stage to analyze changes in neuronal structure and function. B. Example average projection of the transverse imaging plane of CA1 through the microperiscope in a mouse sparsely expressing Thy1-GFP-M. C. Weighted projection (see Methods) of the apical dendrite shown in the dashed box of (B). D. Filtered and binarized image (see Methods) of the dendrite shown in (C), to allow for the identification and classification of individual dendritic spines, and to mask extraneous projections. E. Binarized thresholded projections of a gaussian-averaged dendrite from apical CA1, taken 24 hours apart in the four archetypal stages of estrous. Circles indicate spine addition, triangles indicate spine subtraction, both relative to diestrus. Color indicates spine type (yellow = filopodium, red = thin, blue = stubby, navy = mushroom), with legend representative of the classic morphological features of each type. F. Percent spine density change from baseline (averaged across stages) for each recorded dendrite across consecutive recordings taken at 12 hour intervals. Stage lengths are interpolated to 24 hrs to allow plotting of all dendritic segments (see Methods; D = diestrus, P = proestrus, E = estrus, M = metestrus). Blue lines indicate individual dendritic segment time courses, while the bolded navy line indicates mean spine density ± standard error. Wilcoxon signed-rank test against a grand mean of all dendrites included at each timepoint. \*\**p* < 0.01, \*\*\**p* < 0.001, \*\*\*\**p* < 0.0001. The number of spines of each type (filopodium (yellow), thin (red), stubby (blue), and mushroom (navy)) per 10 µm section of CA1 apical dendritic segment for each estrous stage, mean ± standard error. Bars indicate significant modulation across all stages for thin and stubby spines (linear mixed effects model). \**p* < 0.1, \*\**p* < 0.01.

Early cross-sectional work identified significant estrous-mediated fluctuations in dendritic spine density on the apical dendritic branches of CA1 pyramidal neurons^10–14^. These *ex vivo* studies were instrumental in demonstrating endocrine modulation of synaptic density, but lacked the ability to measure the same synapses across the estrous cycle. *In vivo* hippocampal imaging allows for longitudinal measurement of dendritic spines, however this technique typically employs aspiration of overlying neocortex for a top-down view of hippocampus^38–40^, fundamentally limiting the visibility of apical dendritic processes, which in dorsal CA1 extend ventrally from the soma^41^. To image the transverse plane of the hippocampal circuit *in vivo*, we developed chronically implanted glass microperiscopes, which allowed optical access to the apical dendritic arbor of pyramidal neurons, the principal site of estrous-modulated synaptic remodeling (Fig. 1B)^42^. To longitudinally image dendritic spine morphology without background contamination, we implanted the microperiscope into Thy1-GFP-M mice, which express GFP in a sparse subset of pyramidal neurons (Fig. 1A,B)^42^. To resolve individual spines along the dendrite, images were taken from several axial planes spanning the segment, and a composite image was generated using a weighted average of individual planes (Fig. 1C). We reduced noise by filtering and binarizing, and isolated dendrites of interest for tracking across days (Fig. 1D). We used custom software to automatically detect spines across sessions, and then manually checked identified spines using an additional interactive GUI (fig. S1; see Methods).

Dendritic spines fall into one of four major morphological subtypes: filopodium, thin, stubby, and mushroom^43–45^. We classified dendritic spines into their relative subtypes based on spine shape parameters (Fig. 1E) and evaluated density and turnover as a function of spine type. Consistent with early histological results, spines in the living mouse were primarily added during the high-estradiol stage proestrus, and pruned during the low-estradiol stage estrus (Fig. 1E,F).

Across a population of 21 dendritic segments and n = 6 mice, spine density was significantly modulated by estrous stage. During proestrus we observed an increase in spine density of 11.5% ± 0.2% (*p* < 10^-4^, Wilcoxon rank sum test; mean ± sem), compared to the global mean across stages, while in estrus we observed a decrease in spine density of 12.4% ± 0.2% (*p* < 10^-4^, Wilcoxon rank sum test; mean ± sem; Fig. 1F). Spine density during diestrus and metestrus was not significantly different from the global mean (D: *p* = 0.1284, M: *p* = 0.6450, Wilcoxon rank sum tests; Fig. 1F). The estrous cycle had differential effects on specific spines types, with significant modulation of thin spines (*p* = 0.0019, *F*(3) = 5.41, linear mixed-effects model (lme), fixed effect for stage, random effect for mouse for all comparisons) and modest modulation of stubby spines (*p* = 0.0497, *F*(3) = 2.72, lme); while mushroom spine density remained largely stable across stages (*p* = 0.8232, *F*(3) = 0.30, lme; Fig. 1G). Filopodia made up only 3% of total spines, and did not appear to be significantly modulated by estrous (*p* = 0.1948, *F*(3) = 1.60, lme). We note that the width of filopodia may fall below the functional diffraction limit of our imaging system, and as a result may be undercounted. However, recent results indicate that filopodia are primarily silent synapses, and do not contribute to changes in excitatory neurotransmission^46^. Together, these results indicate that the estrous cycle drives robust changes in the density of intermediate-sized spines that are likely to contribute to functional excitatory input and neuronal processing^46^.

### Morphological spine dynamics are shaped by estrous cycle stage

One advantage of longitudinal monitoring of dendritic spines is that it is possible to address outstanding questions regarding the dynamics of different spine types. For instance, are new spines added during proestrus immediately pruned during estrus, or do they become a stable part of the synaptic milieu? To address this, we first measured the survival fraction of all dendritic spines present on the first recording session that were still present on session *n*^47, 48^. For these spines, after 10 consecutive recordings (i.e. 5 days), 82.2% ± 2.2% (mean ± sem) of spines found on session 1 remained (Fig. 2A), consistent with measurements taken in a mixed sex cohort^42^. We then calculated a survival fraction curve for spines added during proestrus, where the only spines included in the analysis were spines for which diestrus was observed prior to proestrus to confirm spontaneous addition, and when a full cycle of recordings were taken after the spine appeared. After entering estrus 52.7% ± 5.7% (mean ± sem) of spines were immediately pruned, and at the end of 10 recordings only 35.9% ± 3.2% (mean ± sem) of proestrus-added spines remained (Fig. 2A). This indicates that while the majority of spines added during proestrus are lost when estradiol levels drop during estrus, a substantial fraction are incorporated into the total functional synaptic population.

**Figure 2.**
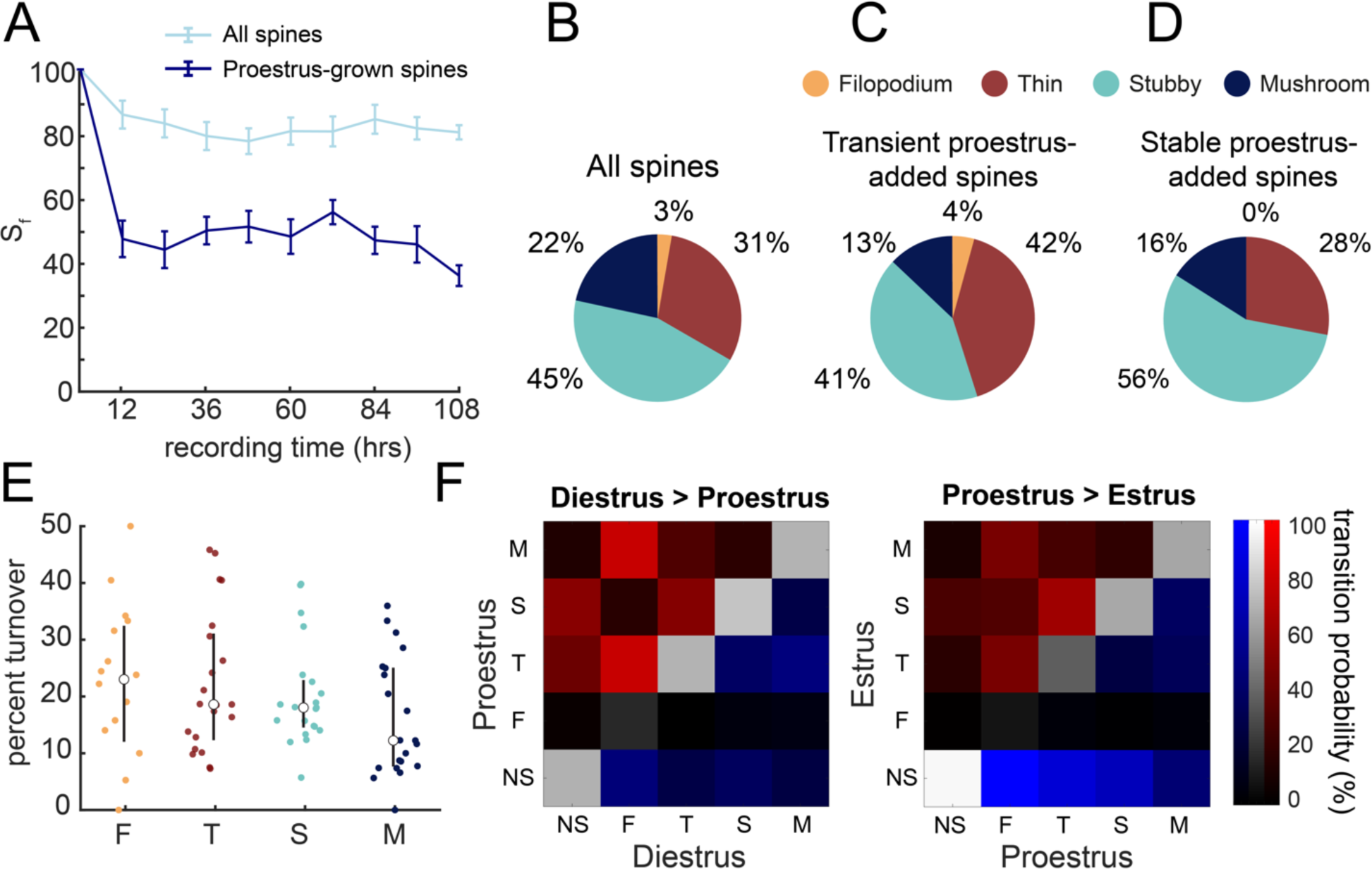
Dendritic spine properties dynamically shift across the estrous cycle. A. Survival fraction of all spines present on recording *n* that were present on recording 1 (at 12 hr intervals) for all spines that were present on the first recording session (light blue), as well as spines that spontaneously appeared during proestrus (navy), where recording 1 is considered the time point during proestrus at which the spine was first observed. Mean ± standard error. B. The proportion of classical spine types across the entire dendritic spine population. Yellow = filopodium, red = thin, blue = stubby, navy = mushroom. C. The proportion of spine types for spines that appeared during proestrus but were immediately pruned in the recording following proestrus (transient spines). D. The proportion of spine types for spines that appear during proestrus and are maintained throughout the entire next cycle (stable spines). E. Percent turnover by session of all spines, analyzed respective to spine type (yellow = filopodium, red = thin, blue = stubby, navy = mushroom). Mean ± standard error. F. Transition matrix for all spines that were present in diestrus and recorded until proestrus (D>P), and spines that were present in proestrus and recorded until estrus (P>E). Spines are classified in order of least to most stable (NS = no spine, F = filopodium, T = thin, S = stubby, and M = mushroom), as established by the survival fraction shown in (E). Transition probability is shown by brightness, where completely dark is 0% transition probability and completely bright is 100% transition probability. Transitions to more stable spine types are pseudocolored in red, persistence within the same type is shown in white, and transitions to less stable types are shown in blue.

Of the spine population classified as “proestrus-added”, we further broke spines down into two categories: those that were pruned at some point in the cycle after they first appeared (“transient spines’’), or those that were maintained throughout the entirety of the cycle after they appeared (“stable spines”). When these categories are broken down by spine type, their classifications are noticeably different from the spine type ratio of the general spine population (2.8% filopodium, 30.6% thin, 45.1% stubby, 21.6% mushroom), and from each other.

Compared to the general spine population, transient spines have 10.3% more spines classified as thin and 8.7% fewer mushroom spines (Fig. 2B,C), while stable spines had 10.9% more stubby spines and 5.6% fewer mushroom spines compared to the general population (Fig. 2B,D). Strikingly, stable spines had 12.9% fewer thin spines and 14.1% more stubby spines than transient spines, suggesting that thin spines were more likely to be pruned than their stubby counterparts (Fig. 2C,D).

When percent turnover, i.e. net change in spines per session, is broken down by spine type, it suggests the following stability hierarchy, from least to most stable: 1) filopodium (23.0% ± 11.1%, mean ± sem), 2) thin (18.6% ± 6.3%, mean ± sem), 3) stubby (18.0% ± 3.5%, mean ± sem), and 4) mushroom (12.2% ± 4.6%, mean ± sem) (Fig. 2E). Transition matrices reveal that spines transition from less stable to more stable types during the transition from diestrus to proestrus (Fig. 2F, left). Conversely, spines are more likely to be pruned (spine > no spine) rather than transitioning to less stable spine types in the transition from proestrus to estrus (Fig. 2F, right). This suggests that, in addition to spinogenesis and spine pruning, there are more subtle estrous-driven morphological changes that may have implications for the functional connectivity of the hippocampal network.

### Estrous modulates apical dendritic activity

Given the significant changes in spine turnover observed across estrous stages, we next investigated estrous modulation of dendritic activity in CA1. To accomplish this, we used a Cre-dependent GCaMP6s transgenic mouse line (TIT2L-GCaMP6s) injected with diluted CaMKIIα- Cre virus, resulting in sparse and stable GCaMP6s expression in CA1 pyramidal neurons with low neuropil contamination (Fig. 3A-C; movie S1). Using this combinatorial approach, we quantified changes in local calcium responses as a measure of postsynaptic activity in CA1 dendrites. To target apical dendrites, which are the major site of estrous-mediated spine turnover, mice were implanted with glass microprisms that allowed optical access to the somatodendritic axis of CA1 neurons (Fig. 3A,C)^42^.

**Figure 3.**
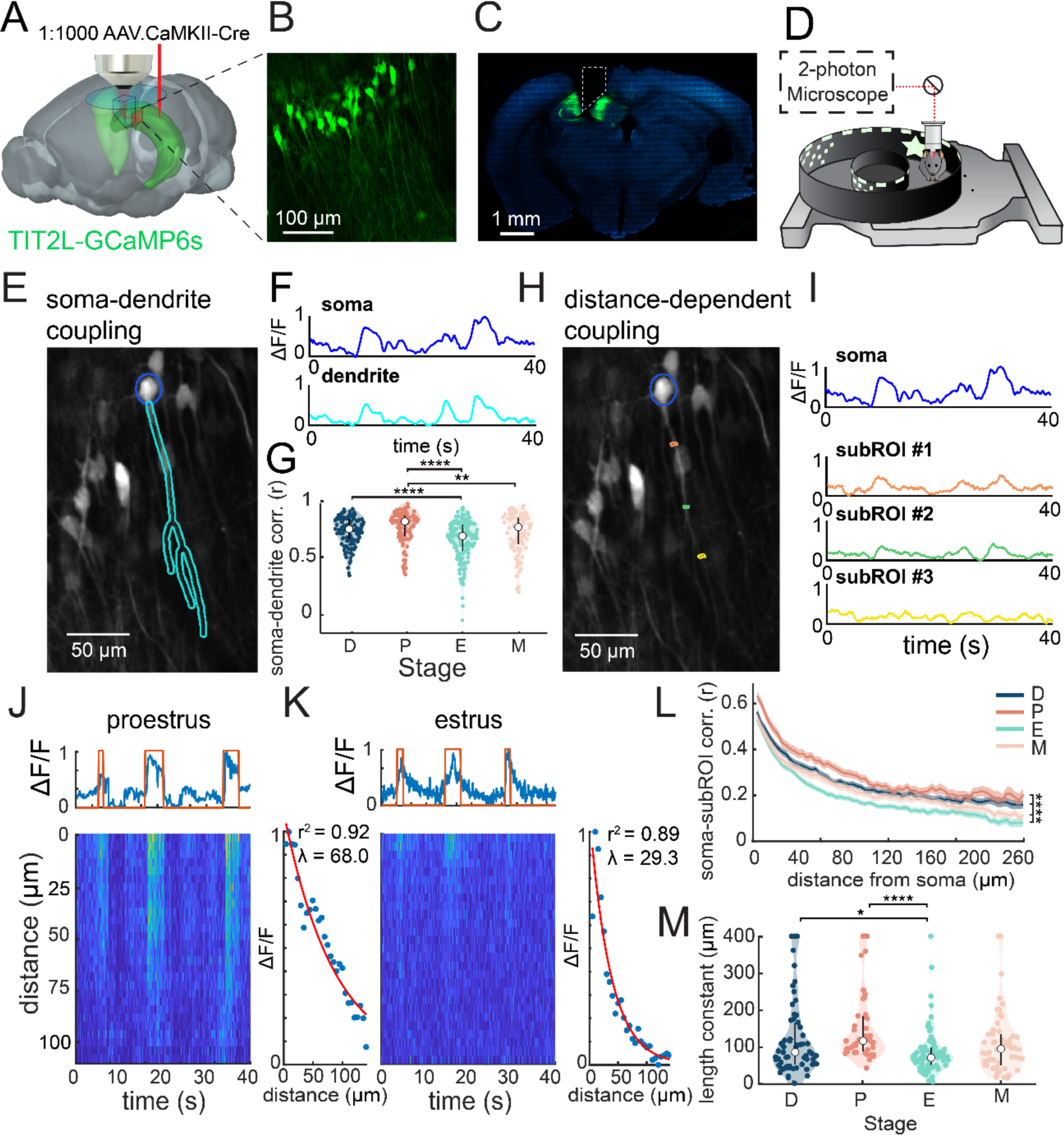
Estrous modulates somatodendritic coupling and action potential propagation. A. Schematic showing microinjection of dilute CaMKII-Cre AAV into TIT2L-GCaMP6s transgenic mouse for sparse, stable dendritic expression prior to implantation of hippocampal microperiscope. B. Average projection of sparse CA1 neurons expressing GCaMP6s throughout the somatodendritic axis, as viewed through the microperiscope. C. Confocal image of a TIT2L-GCaMP6s mouse with viral injection of dilute CaMKII-Cre into CA1 in a coronal slice from a mouse implanted with a microperiscope (boundaries of implantation shown in dotted box). D. Schematic of the air-lifted carbon fiber floating track that the mice explored during imaging. Average projection of an example place cell showing somatic and apical dendritic ROI selection, from which ΔF/F is extracted. E. Average projection of an example pyramidal cell in CA1 showing soma and apical dendrite ROI selection. F. Somatic and apical dendritic ΔF/F across a 40 second interval as the animal traverses a floating chamber. Dendritic ΔF/F (cyan) is normalized to somatic ΔF/F (blue). G. Correlations between somatic and dendritic ΔF/F from the entire recording across estrous stages. Significance between correlations across stages given by pairwise linear mixed effect models. Mean ± standard error. \*\**p* < 0.01, \*\*\*\**p* < 0.0001. H. Average projection of an example pyramidal cell in CA1 showing somatic and subROI selection, with three highlighted 6 µm-wide subROIs ∼50 µm apart along the dendritic segment. I. Sub-ROI ΔF/F at increasingly distal points along the main apical dendritic branch (orange = most proximal, green = medial, yellow = most distal), normalized to somatic signal (top, blue). J. Example pyramidal cell response from a recording taken in proestrus. Forty seconds of normalized somatic ΔF/F (blue, top) is shown, aligned with a logical trace (red, top) set to true during frames when a bAP occurred. A heatmap shows the response of each subROI over the same 40 second epoch. The subROI response during each bAP is normalized to mean somatic ΔF/F during the bAP, then all bAP responses are averaged to create a representative bAP curve, which is fit to an exponential to determine the length constant (right). K. Example pyramidal cell response from a recording taken in estrus. Somatic signal, subROI signal, and bAP fitting is shown as described in J. L. Correlations between soma and increasingly distal subROIs along the dendritic branch across estrous stages. Bar indicates significant modulation across all stages (linear mixed effects model). Mean ± standard error. \*\*\*\**p* < 0.0001. Distributions of the average length constant from all CA1 pyramidal cells analyzed across estrous stages. Pairwise linear mixed effect model. Mean ± standard error. \**p* < 0.1, \*\*\*\**p* < 0.0001.

To measure activity in CA1 pyramidal cells across the estrous cycle, two-photon imaging was performed at 12-hour time points. Since the two-photon imaging requires head fixation, mice ran on an air-floated platform, allowing them to explore a circular track lined with phosphorescent visual landmarks (Fig. 3D; movie S2). The experiment was otherwise conducted in darkness in an enclosed light box in order to eliminate the influence of distal visual cues outside the floating chamber.

The increased spine density observed during proestrus is thought to contribute to a net increase in dendritic excitation, which may in turn increase correlations between somatic and dendritic activity due to greater intrinsic excitability^49–51^. Our recordings revealed significantly increased soma-dendrite coupling during the proestrus stage, and significantly decreased soma-dendrite coupling during the estrus stage (*p* < 10^-4^, *F*(3) = 10.37, lme, fixed effects for stage, random effects for mouse; Fig. 3E-G; table S2).

We next asked what specific aspects of apical dendritic activity were leading to estrous-dependent differences in somatodendritic coupling. To accomplish this, we subdivided dendritic ROIs into 6-μm segments (“sub-ROIs”) and re-extracted, processed, and analyzed sub-ROI signals (Fig. 3H,I)^49^. All stages showed a monotonic decrease in correlation between sub-dendritic ΔF/F and somatic ΔF/F as a function of distance along the dendrite, however this decrease was significantly more pronounced in estrus compared to proestrus (*p* < 10^-4^, *F*(3) = 212.26, lme, fixed effects for stage, random effects for mouse; Fig. 3L; table S2).

Backpropogating action potentials (bAPs) are known to play an important role in regenerative dendritic events^52–54^ and behavioral time scale plasticity^25, 55, 56^. The variable somatodendritic coupling seen during particular estrous stages could potentially influence the spread of bAPs into dendritic arbors and modulate CA1 plasticity. To test this possibility, bAPs were identified in somatic ΔF/F and aligned with dendritic responses. Then, within-cell bAPs were averaged, normalized to the somatic response, and fit with an exponential decay function to determine the bAP length constant for each cell (Fig. 3J,K; See Methods). When analyzed by estrous stage, length constants were significantly higher in proestrus, when somatodendritic correlations are high, compared to estrus, when somatodendritic correlations are low (*p* < 10^-4^, *F*(3) = 6.55, lme, fixed effects for stage, random effects for mouse; Fig. 3M; table S2). To determine the extent to which spatial coding may be modulated by estrous stage, these analyses were recapitulated only considering place cells, which fired consistently in the same location along the circular track. When only place cells were analyzed, the results remained consistent, with significantly greater somatodendritic coupling, soma-subROI coherence over distance, and bAP spread during the high-estradiol stage proestrus (fig. S2; table S4). This suggests a postsynaptic mechanism for estrous-modulated somatodendritic coupling, in which changes in synaptic density lead to differences in excitatory signal propagation.

### Place cells measured in floating environments exhibit stable responses and coordinated remapping

Does estrous cycle modulation of dendritic activity influence place cell activity in CA1? To address this, we imaged large populations of place cells across the estrous cycle in actively navigating head-fixed animals. Although virtual reality has been used in studies of spatial memory, it remains unclear how similar rodent hippocampal activity in real-world environments is to that in virtual reality^57, 58^. In fact, two-dimensional place fields are limited in virtual reality, with impaired spatial tuning and theta rhythm frequency^57–59^. To this end, we used a floating chamber with multiple sensory modalities to drive reliable place cell activity across laps, and remapping between different environments^59^. To maximize the population of cells in the imaging field, we used surgically implanted glass cylinders and imaged CA1 somata (Fig. 4A,B; fig. S3A; movie S3). This was necessary to capture a sufficient population of place cells for analysis, which prior studies suggest make up only 20-30% of the total population of mouse CA1 neurons^31, 42, 60^.

**Figure 4.**
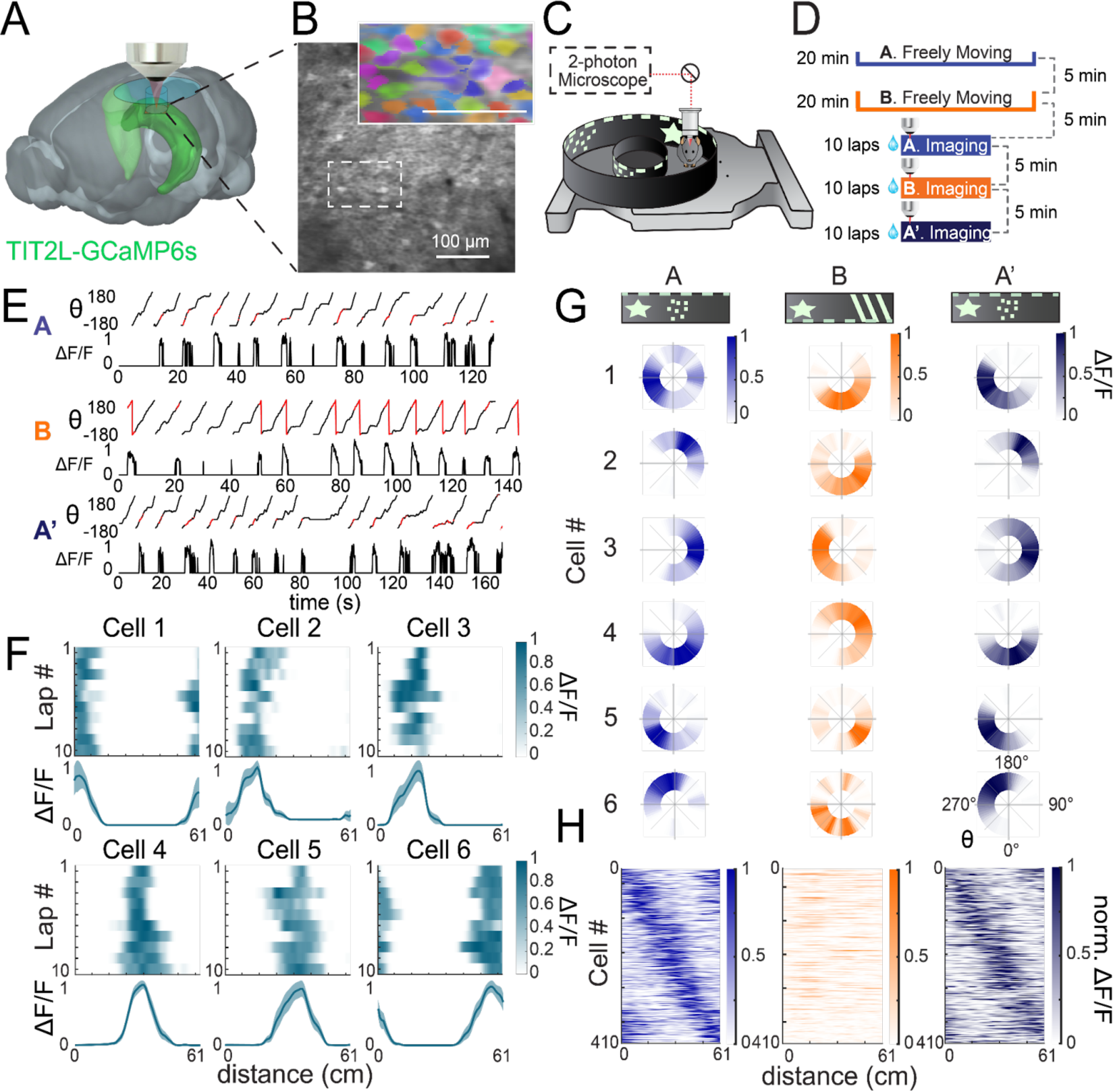
Changes in environmental cues induce remapping in CA1 place cells. A. Three-dimensional schematic of a plug implant for top-down two-photon calcium imaging of CA1 cell bodies in CaMKIIa-Cre x TIT2L-GCaMP6s mice. B. Average projection of CA1 somata through the glass plug implant, scale bar = 100 µm. Inset, ROI selection from the region shown in dashed box, with randomized color scheme. C. Schematic of floating chamber experimental design, where the mouse is head-fixed for two-photon imaging. The walls of the chamber are lined with fluorescent local visual cues and the base of the chamber includes interchangeable textural cues. D. Experimental design schematic, where an open-top box represents a freely moving mouse during the acclimation period, and a closed box represents a head-fixed mouse during the recording period. Mice are allowed to run 10 laps, motivated by water reward, before being moved to the resting cage and subsequently introduced to the next environment. E. Angular position output of the floating chamber tracking system aligned with ΔF/F transients of a single CA1 place cell across three environments : A, B, and A’. ΔF/F transients are shown in red in the angular ΔF/F trace for visualization. F. Smoothed ΔF/F transients from six example cells across laps around the floating chamber (top), as well as lap-averaged one-dimensional tuning curves (bottom; mean ± sem). G. Single-cell angular ΔF/F from six example cells averaged across laps as the mouse moves from environment A (top left) to environment B (top center), and back to environment A (A’; top right). A = blue, B = orange, A’ = purple. Note similar place fields in environment A and A’, but shifted place field location in environment B. Place cell responses in CA1 from an example recording across the three environments: A (blue), B (orange), A’ (purple). Average responses (normalized ΔF/F) of all place cells are ordered by peak position in environment A for all three environments. Responses in environment A are cross validated by determining peak position in odd trials and plotting even trials. Note that place fields are in similar positions for environment A and A’, but remap in environment B.

Since previous studies have shown that sensitivity to environmental context does not reach a peak until the mouse has been exposed to a familiar environment for at least 21 days^61, 62^, the mice were acclimated to the floating chamber for 21 days prior to recording, including head fixation and water reward (fig. S3B). Different environments in the floating chamber were defined by distinct patterns of phosphorescent visual cues (A: dots, B: stripes) and textural floor cues (A: foam, B: bubble wrap; Fig. 4C), and experiments were carried out in a dark chamber to reduce the influence of distal cues. To aid in establishing a spatial representation of the floating chamber as a repeating circular track, and to give a reference point for remapping, one visual cue was kept constant between environments and aligned to a static textural “start” point, marked by a divot in the base of the chamber. When the mouse completed a lap and returned to the start point, they received a water reward. Prior to recording sessions, mice were allowed to freely explore each environment for 20 minutes, with a 5-minute interlude in their home cage in between exposures (Fig. 4D). During imaging, the mice were allowed to run ten consecutive laps, then were placed back in the home cage for a rest interval of 5 minutes before introduction to environment B. This was repeated for the transition from environment B back to environment A (A > B > A’) (Fig. 4D).

The position of the floating chamber was measured using a magnetic tracking system and aligned to neural fluorescence traces (ΔF/F), for which calcium transients were extracted and smoothed while all other points were masked to zero, to correct for slow changes in fluorescence (Fig. 4E; fig. S2B)^38^. Place cells had well-defined fields of elevated neuronal activity that were stable across laps at particular locations in the circular track (Fig. 4E,F). When averaged across laps for each of the three environments (A > B > A’), the preferred angular position of place cells consistently shifted from environment A to environment B, then returned to their original position when reintroduced to environment A (A’; Fig. 4G). At the population level, CA1 place cells tiled the length of the circular track, remapping from environment A to B, but remaining stable when reintroduced to environment A (A’; Fig. 4H). Despite the track maintaining identical geometry and dimensions between environments, these results demonstrate that changes to visual and tactile cues provided sufficient sensory context to induce global remapping in the CA1 place cell population^59^.

### Place cell stability is modulated by estrous cycle stage

Having established strong place cell responses in the floating chamber, we next measured the stability of spatial representations within the same environment and the flexible remapping of spatial responses between distinct environments as a function of estrous stage. Previous electrophysiological studies have demonstrated that the basic firing properties of place cells are stable across the estrous cycle^63^. Consistent with these results, we found that place field width (*p* = 0.1375, *F*(3) = 1.36, lme; FWHM), spatial information (*p* = 0.2457, *F*(3) = 0.92, lme; bits/inferred spike), and mean event rate (*p* = 0.1196, *F*(3) = 1.13, lme) were not significantly modulated by estrous cycle stage (fig. S4A-C; table S5). Estrus did show significantly lower lap-wise stability than either proestrus (*p* < 10^-4^, *F*(1) = 54.62, pairwise lme) or metestrus (*p* = 0.0014, *F*(1) = 13.58, pairwise lme), however the stability of cells in estrus is still sufficient to suggest the formation of strong place fields (fig. S4D). Despite consistency in basic firing properties, when our results were analyzed as a function of stability across the same environment (A> A’) and remapping between different environments (A > B), we found significant estrous modulation of spatial representations. As exemplified by single-cell traces, neurons in proestrus exhibited greater stability in spatial tuning across the same environments (Fig. 5A). When sorted by latency in environment A, the place cell population response visibly preserves more similar place fields from environment A to A’ in proestrus compared to estrus (Fig. 5B).

**Figure 5.**
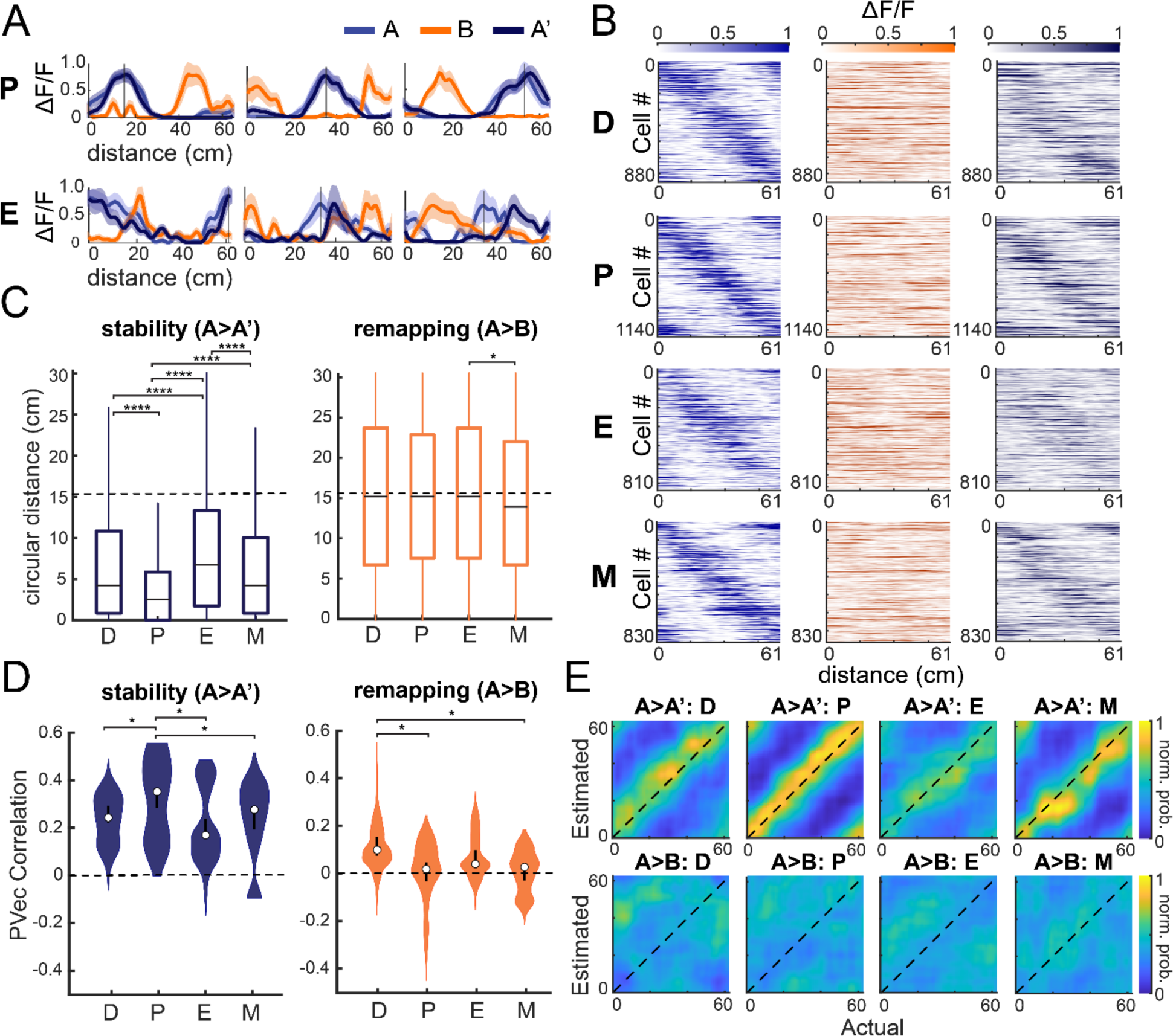
The stability of place fields is modulated by estrous cycle stage. A. Six example cells showing place fields averaged across laps (mean ± standard error) in the proestrus (P) and estrus (E) stages. Positional tuning curves for environment A shown in blue, B in orange, and A’ in purple. Note that neurons imaged during proestrus exhibit more stable place fields between environments A and A’ than neurons imaged during estrus. B. Cross-validated average responses (normalized ΔF/F) of all place cells ordered by peak position in environment A in each estrous stage. D = diestrus, P = proestrus, E = estrus, M = metestrus. C. Distribution of the circular distance between place cell peaks across environments (either A to A’, i.e., stability, or A to B, i.e., remapping) in each estrous stage. Box plots show mean (horizontal line), first and third quartile (box), and maximum and minimum (whiskers). A dotted line at zero is shown for reference. Pairwise linear mixed effects model. \*\*\**p* < 0.001, \*\**p* < 0.01. D. Population vector correlation distributions across estrous stages in the same environment A (stability; A>A’) and different environments (remapping; A>B). Distributions were created by bootstrapping data from n = 6 mice and n = 3679 cells over 100 iterations, sampling with replacement. A dotted line at zero is shown for reference. Overlay indicates mean ± bootstrapped 95% CI. Asterisks denote significance. Significance was defined as lower 5% CI higher than higher 95% CI. Probability density plot of the average prediction accuracy from decoders predicting position in environment A’ (A>A’, top) compared to environment B (A>B, bottom). Responses for each of the four estrous stages are averaged across one recording per stage for each of n = 6 mice.

To quantify this phenomenon, we first calculated the circular difference in place cell peak between environments A>A’ (stability; left), and A>B (remapping; right). From environments A to A’ stability was significantly greater in proestrus and lower in estrus (*p* < 10^-4^, *F*(3) = 105.46, lme, fixed effects for stage, random effects for mouse; Fig. 5C). We found only minor differences in single-cell responses in the remapping condition (*p* = 0.0427, *F*(3) = 2.73, lme, fixed effects for stage, random effects for mouse; Fig. 5C). To confirm population-level effects, we calculated the population vector (PVec) correlation to assess the degree of stability and remapping across environments and stages in the CA1 place cell population^64, 65^. A PVec correlation of 0 indicates strong remapping, whereas a PVec correlation of 1 indicates complete stability (i.e., no remapping at all)^64, 65^. Consistent with our single-cell analysis, we found that place cells in proestrus exhibit significantly greater PVec correlations (i.e. more stability), while place cells in estrus exhibit significantly lower PVec correlations (i.e. less stability), as shown by comparison of the bootstrapped 95% confidence intervals (Fig. 5D). Also consistent with our single-cell analysis, differences in remapping across stages showed only minor differences.

Despite the observed changes in stability and remapping, we did not find estrous-dependent effects on total moving time, speed, or anticipatory reward behavior (fig. S3C; fig. S5; table S6).

Finally, from the activity of these neurons, we trained a linear decoder on place cell population activity to predict the animal’s location along the circular track^65, 66^. After training the decoder on place cell responses in environment A, the decoder was asked to predict location when the animal was in a different environment (B) versus in the same environment (A’; Fig. 5E). As expected, the decoded position was much closer to the actual position in environment A’, in which place fields were stable, than in environment B, in which place cells had undergone global remapping (Fig. 5E). Consistent with single-cell and PVec analysis, proestrus also showed significantly lower decoder error than estrus when estimating position from the same environment, as shown by non-overlapping 95% CI (fig. S6). The performance of this decoder supports the result that the high-estradiol stage, proestrus, exhibits significantly more stability between environments than the low-estradiol stage estrus. Taken together, these results are the first to demonstrate that spatial representations in the hippocampus are modulated by cyclic endocrine factors, from the synaptic to population level. This underscores a growing body of work suggesting that consideration of hormonal signaling is critical to understanding the intrinsic dynamics of spatial coding in the brain.

## Discussion

Here we investigated the role of the murine estrous cycle on hippocampal structural and functional plasticity at the synaptic, cellular, and population levels. Building on previous cross-sectional studies, we found that dendritic spine density is fundamentally shaped by the estrous cycle, with a subset of proestrus-added spines becoming part of the stable synaptic milieu (Figure 1, 2)^10–14^. The dendritic activity of CA1 place cells was also estrous-modulated, showing greater infiltration of bAPs into the apical dendrite during periods of high estradiol, and higher coupling of somatodendritic activity (Figure 3). Finally, we also found that population-level spatial coding was significantly more stable across exposures to the same environment (A > A’) during the high estradiol stage, proestrus, and significantly less stable during the low estradiol stage, estrus (Figure 5). These results are the first to demonstrate *in vivo* functional modulation of hippocampal circuit function by the estrous cycle, offering a window into the role of cyclic plasticity in shaping mammalian spatial cognition.

Recent work has led to new conceptions of the role of synapse-level plasticity in the formation and stability of spatial representations. These theories of place field formation hypothesize that CA1 place fields are driven by local activity in the dendritic branches of hippocampal neurons, in the form of bAPs, plateau potentials, and NMDA spikes^25^. Such plasticity models propose that bAPs underlie non-Hebbian plasticity mechanisms shown to rapidly initiate spatial coding in CA1 neurons and stabilize existing place fields^25, 55, 56^. This suggests that the greater bAP spread observed in proestrus may contribute to the population-level stability of place cells^25^. Moreover, several computational models of place cell formation demonstrate that increased excitatory dendritic input leads to greater place field stability, consistent with our findings that periods of higher synaptic density exhibit more stable place fields within the same environment^66–68^.

These models raise the possibility that local plasticity in downstream areas may demonstrate cyclic functional modulation by circulating hormones. While estrous-dependent changes in dendritic spine density have not been observed in medial entorhinal cortex or somatosensory cortex^69^, cross-sectional synaptic fluctuations similar to those observed in the hippocampus have been observed in ventromedial hypothalamus^70^, prefrontal cortex^71^, and amygdala^72, 73^. Further studies investigating the role of endocrine factors in these areas will be critical for developing a more complete understanding of hormone modulation in the brain.

Future studies will also be instrumental in uncovering the molecular mechanisms underlying estrous modulation of structural and functional hippocampal plasticity. Classical estrogen receptors ERα and ERβ play a role in the formation of functional synapses in response to exogenous estradiol administration^1, 15, 22^. However, it remains unclear to what extent these receptors modulate the function of whole cells or networks, and the role that G-protein coupled estradiol receptors or progesterone receptors play in estrous-dependent changes in hippocampal morphology and function. Advances in CRISPR^74^, shRNA^75^, and transgenic technology^76^ may pave the way for a more complete understanding of the mechanisms underlying changes in spatial coding across the estrous cycle.

Given our findings, an important question to address is: what are the evolutionary mechanisms underlying such significant cyclic changes to the hippocampal network? The estrous cycle is fundamental for reproduction, with estradiol levels peaking during proestrus immediately prior to ovulation in estrus, when estradiol levels plummet up to 15-fold^34, 77^.

Previous studies suggest that wandering behavior increases in late proestrus, which may be a mechanism for timing copulation during the ovulatory period^33, 78^. Some studies suggest that navigation behavior in rodents switches from an allocentric to an egocentric strategy in the transition from proestrus to estrus, which supports our findings that place cell stability in a familiar environment is greater during proestrus^79–81^. An important qualification to this interpretation is that recent deep-learning interrogations of mouse behavior across the estrous cycle concluded that spontaneous motor behavior is not modulated by the estrous cycle, and that the individual identity of the animal is significantly easier to decode than estrous stage^82^. Additionally, studies investigating the role of estradiol in completing navigation tasks like the Morris water maze and radial arm maze demonstrate contradictory results depending on a range of methodological factors, with different studies showing either enhanced or impaired spatial memory after estradiol administration^1, 83–86^. Therefore, additional studies will be required to fully understand the impact of cyclic hippocampal modulation on spatial memory and behavior, and to elucidate any evolutionary logic underlying hippocampal changes across the estrous cycle.

It is also important to note that the estrous cycle in rodents is often linked to the erroneous perception that female animals are more variable than males, resulting in the chronic underrepresentation of female animals in research studies^87, 88^. Several studies to date, however, definitively demonstrate that male rodents actually display greater behavioral variability than females^82, 89^. Furthermore, estradiol exerts a strong modulatory influence on the male hippocampus, where it is present at higher levels than in the female hippocampus^1, 90^ and, as in females^15^, induces synaptic in hippocampal neurons^91, 92^. The presence of a hormonal cycle is also not limited to female animals. For example, circadian cycles are common in mammalian species, with steroid hormone levels typically peaking in the morning and decreasing in the evening^93^. Male mice additionally exhibit several cyclic peaks of testosterone, progesterone, and cortisol throughout the day^94^. In contrast to the 4-5 day cycle of female mice, the rapid time course of male hormone fluctuations make it difficult to capture cyclic plasticity changes, and vaginal cytology is not an option for non-invasive staging of hormonal state.

However, with the rise of chronic electrophysiological recordings^95, 96^ and new technologies for plasma hormone assessment^97, 98^, future studies could investigate the effects of male hormonal modulation on hippocampal circuitry.

Taken together, our findings demonstrate that naturally cycling endocrine factors are robust modulators of spatial memory circuits, shaping hippocampal structure and function on a previously unprecedented scale.

## Materials and Methods

### Animals

For dendritic morphology experiments, Thy1-GFP-M (Jax Stock #007788) female transgenic mice (*n* = 6, 9-17 week old females) were used for sparse expression of GFP throughout the forebrain. For sparse dendritic calcium imaging, TITL2-GC6s-ICL-TTA2 (Jax Stock #031562) transgenic mice were injected with a CaMKIIα-Cre virus (Addgene #105558-AAV1) diluted 1:10,000 in CA1 (*n* = 5, 14-28 week old females). For forebrain-wide calcium indicator expression, CaMKIIα-Cre (Jax Stock #005359) × TITL2-GC6s-ICL-TTA2 (Jax Stock #031562) double transgenic mice (*n =* 6, 8-12 week old females) were bred to express GCaMP6s in excitatory neurons. For imaging experiments, female mice were implanted with a head plate and cranial window and imaged starting 2 weeks after recovery from surgical procedures. The animals were housed on a 12 hr light/dark cycle in cages of up to five animals before the implants, and individually after the implants. All animal procedures were approved by the Institutional Animal Care and Use Committee at University of California, Santa Barbara, CA.

### Estrous Cycle Staging

All control and experimental mice were regularly staged by vaginal lavage starting two weeks post-surgery (once per day prior to imaging and twice per day during imaging). Samples were collected during the light phase of the cycle using 50 µl sterile saline pipetted into the vaginal opening and aspirated several times to obtain a sufficient cell count. The sample was pipetted onto a gel subbed microscope slide and allowed to dry 24 h before staining with Shorr Stain (Sigma Aldrich). Gel subbing was performed in house using standard IHC protocol to coat glass slides in gelatin/CrK(SO4)2 solution^99^. Dried and stained slides were imaged under a compound brightfield OMAX microscope using ToupView software at 10X magnification. Images were first assessed visually, then fed into EstrousNet for automated estrous stage classification^36^. Staging was performed by separate experimenters blind to experimental condition.

### Surgical procedures

All surgeries were conducted under isoflurane anesthesia (3.5% induction, 1.5–2.5% maintenance). Prior to incision, the scalp was infiltrated with lidocaine (5 mg kg^−1^, subcutaneous) for analgesia and meloxicam (2 mg kg^−1^, subcutaneous) was administered preoperatively to reduce inflammation. Once anesthetized, the scalp overlying the dorsal skull was sanitized and removed. The periosteum was removed with a scalpel and the skull was abraded with a drill burr to improve adhesion of dental acrylic.

For transverse hippocampal imaging (Figures 1–3), we used a custom-designed glass microperiscope (Tower Optical) consisting of a 1 × 1 × 1 mm^3^ square base and a 1 mm right angle prism, for a total length of 2 mm on the longest side. The hypotenuse of the right angle prisms were coated with enhanced aluminum for internal reflectance. The microperiscope was attached to a 5 mm diameter coverglass (Warner Instruments) with a UV-cured optical adhesive (Norland, NOA61). Prior to implantation, the skull was soaked in sterile saline and the cortical vasculature was inspected to ensure that no major blood vessels crossed the incision site. If the cortical vasculature was suitable, a 3–4 mm craniotomy was made over the implantation site (centered at 2.0 mm posterior, 1.2 mm lateral to Bregma). For implantation, a 1 mm length anterior-to-posterior incision centered at –2.1 mm posterior, 1.2 mm lateral to Bregma was then made through the dura, cortex, and septal (mediodorsal) tip of the hippocampus to a depth of 2.2 mm from the pial surface with a sterilized diamond micro knife (Fine Science Tools, #10100- 30) mounted on a manipulator. Care was taken not to sever any major cortical blood vessels. Gelfoam (VWR) soaked in sterile saline was used to remove any blood from the incision site. Once the incision site was free of bleeding, the craniotomy was submerged in cold sterile saline, and the microperiscope was lowered into the cortex using a manipulator, with the imaging face of the microperiscope facing lateral. Once the microperiscope assembly was completely lowered through the incision until the coverglass was flush with the skull, the edges of the window were sealed with silicon elastomer (Kwik-Sil, World Precision Instruments), then with dental acrylic (C&B-Metabond, Parkell) mixed with black ink. Care was taken that the dental cement did not protrude substantially over the window, as it could potentially scratch the objective lens surface. Given the working distance of the objective used in this study (3 mm), the microperiscope implant enabled imaging from 2250–2600 μm below the coverglass surface, corresponding to approximately 150–500 μm into the lateral hippocampus (the 150 μm of tissue nearest to the microperiscope face was not used for imaging).

For imaging CA1 soma populations (Figures 4–5), we used a custom-designed glass cylinder (Tower Optical) measuring 1.4 mm in diameter, with a length of 1.4 mm. The cylinder was attached to a 4 mm diameter coverglass (Warner Instruments) with a UV-cured optical adhesive (Norland, NOA61). Prior to implantation, the skull was soaked in sterile saline and the cortical vasculature was inspected to ensure that no major blood vessels crossed the implantation site. If the cortical vasculature was suitable, a 3 mm craniotomy was made over the implantation site (centered at 2.0 mm posterior, 1.8 mm lateral to Bregma). For implantation, the dura was removed from over the implant region using fine forceps (FST), and the tissue immediately over the hippocampus was removed with a 1.5 mm diameter sterile biopsy punch, which was was lowered 1 mm into the tissue using a micromanipulator, centered at the implantation site and tilted 10 degrees laterally. Gelfoam (VWR) soaked in sterile saline was used to remove any blood from the incision site. Once the incision site was free of bleeding, the craniotomy was submerged in cold sterile saline, and the glass cylinder was lowered into the incision using a micromanipulator until the bottom surface was flush with the dorsal surface of the hippocampus. The implant was tilted 10 degrees laterally to conform to the surface of the hippocampus. Once the implant was placed, the edges of the window were sealed with silicon elastomer (Kwik-Sil, World Precision Instruments), then with dental acrylic (C&B-Metabond, Parkell) mixed with black ink. For these experiments, the imaging plane was approximately 150–200 μm below the bottom of the cylinder to allow imaging of the stratum pyramidale of subregion CA1.

After implantation of the glass microperiscope or cylinder, a custom-designed stainless steel head plate (https://www.emachineshop.com/) was affixed using dental acrylic (C&B-Metabond, Parkell) mixed with black ink. After surgery, mice were administered carprofen (5–10 mg kg^−1^, oral) every 24 hr for 3 days to reduce inflammation. Microperiscope designs and head fixation hardware are available on our institutional lab website (https://goard.mcdb.ucsb.edu/resources).

### Floating chamber setup, training, and recording

For measurement of spatial responses, mice were head-fixed in a floating carbon fiber chamber (*100*; Mobile Homecage, NeuroTar, Ltd). The chamber base was embedded with magnets to allow continual tracking of the position and angular displacement of the chamber. Behavioral data were collected via the Mobile HomeCage motion tracking software (NeuroTar, versions 2.2.014, 2.2.1.0, and 3.1.5.2). During imaging experiments, image acquisition was triggered using a TTL pulse from the behavioral software to synchronize the timestamps from the 2P imaging and chamber tracking.

A custom carbon fiber arena (250 mm diameter) was lined with detachable 7 mil waterproof paper (TerraSlate) on the outside of the chamber, as well as lining a removable inner circle (14 cm in diameter and 4.2 cm tall) on the inside of the chamber, to create a circular track. The resulting circumference, along the middle of the circular track, is 61.26 cm. Visual cues on both the outer and inner walls were made with photoluminescent tape (Lockport), and the experiments were carried out in near darkness, reducing influence from distal cues outside of the chamber^59^. Tracks were designed with two sets of cues (A, B) to stimulate place cell activity, with one cue used in both sets at the beginning of the track (star) to enable tests of remapping between environments. Cues were charged for 30 minutes under UV light then allowed to rest for 30 minutes to reach a steady phosphorescent intensity before beginning recordings. The first set of cues were paired with a circular foam base insert (NeuroTar), and the second set of cues were paired with a circular base made from perforated bubble wrap. During homecage rest intervals in between imaging sessions, the experimenter scrubbed the chamber with Rescue, a veterinary-grade disinfectant and deodorizer, to eliminate the influence of odor cues between environments.

After water restriction to 85% of initial bodyweight, mice were acclimated to the arena by the following steps (fig. S3B): (1) On the first day the mice were placed into the chamber with cues for environment A and allowed to freely explore without head fixation for 10 min. Small pieces of hydrogel were scattered along the track to encourage exploration. A piece of plexiglass with holes drilled through was placed on top of the arena to keep the mice from climbing out. They were then returned to their home cage for 5 minutes while environment B was prepared, then these steps were repeated for environment B. This was repeated for 3 days with exposures to each environment increasing by 5 minutes on each consecutive day. (2) On the fourth day, the mice were head-fixed to a crossbar extending over the floating chamber (Figure 4C) and allowed to freely explore the floating chamber for 5 min with water reward given at the completion of each lap. Air flow (2–5.5 psi) was adjusted to maximize steady walking/running.

The mice were then placed back in their home cage before being re-head fixed in environment B and allowed to explore for another 5 minutes. For the next 3 days the head fixation time was increased by increments of 2 min in each environment, as long as the mice showed increased distance walked and percent time moving. (3) On day 8, the mice began the complete head fixation sequence of three environments (A > B > A’), where they were allowed to explore each environment for 5 minutes with a 5 minute rest period in the home cage between each exposure. The exposure period increased by 5 minutes every four days until the mice were comfortably running for 15 minutes in each environment on day 19. (6) On the last two days of the training period (days 20 and 21), mice were acclimated to the full trial setup, in which they were first allowed to freely explore each environment for 20 minutes before being head-fixed and placed on the 2P microscope to allow habituation to the microscope noise, and for each environment were only allowed to run 10 laps under a speed threshold of 200 mm/s (laps over the speed threshold were not counted towards the total number of laps run). This was to account for overexertion during exposure to the first environment, and prevent tiring by the third environment. (5) On day 22, after mice were fully habituated, recording sessions on the 2P microscope were performed every 12 hours for 5 days to catch each stage of the estrous cycle.

Custom software was written to process the behavioral data output by the Mobile HomeCage motion tracking software. Because the Mobile HomeCage motion tracking software sampling rate was faster than the frame rate of our 2P imaging, all behavioral variables (speed, location, polar coordinates, and heading) that were captured within the acquisition of a single 2P frame were grouped together and their median value was used in future analysis. For the polar angle (used to determine the location bin of the mouse along the 1D track), the median was computed using an open source circular statistics toolbox (CircStat 2012a) written for MATLAB^101^. We removed any time points when the mouse was not moving, as is standard for measurement of place fields^39^. This helps separate processes that are related to navigation from those that are related to resting state. To do this, we smoothed the measured instantaneous speed and kept time periods > 1 s that had speeds greater than 20 mm/s (adding an additional 0.5 s buffer on either side of each time period).

### Two-photon imaging

After recovery from surgery and behavioral acclimation, GFP or GCaMP6s fluorescence was imaged using an Investigator 2P microscopy system with a resonant galvo scanning module (Bruker). For fluorescence excitation, we used a Ti:Sapphire laser (Mai-Tai eHP, Newport) with dispersion compensation (Deep See, Newport) tuned to *λ*=920 nm. Laser power ranged from 40 to 80 mW at the sample depending on GCaMP6s expression levels. Photobleaching was minimal (<1% min–1) for all laser powers used. For collection, we used GaAsP photomultiplier tubes (H10770PA-40, Hamamatsu). A custom stainless-steel light blocker (eMachineshop) was mounted to the head plate and interlocked with a tube around the objective to prevent light from the environment from reaching the photomultiplier tubes. For imaging, we used a 16×/0.8 NA microscope objective (Nikon) to collect 760 × 760 pixel frames with field sizes ranging from 829 × 829 μm (1x zoom) to 51.8 × 51.8 μm (16x zoom). Images were collected at 20 Hz and stored at 10 Hz, averaging two scans for each image to reduce noise.

For longitudinal imaging of dendritic structure and activity, imaging fields on a given recording session were aligned based on the average projection from a reference session, guided by stable structural landmarks such as specific neurons and dendrites. Physical controls were used to ensure precise placement of the head plate, and data acquisition settings were kept consistent across sessions. Images were collected every 12 hours for 4.5-5 days in calcium imaging experiments, and 8-11 days in structural imaging experiments, with length of session dependent on the periodicity of the estrous cycle, which here ranged from 4 to 5.5 days.

In all two-photon experiments, imaging took place every 12 hours, typically beginning at 6am and 6pm. Imaging periods were conducted in darkness out of necessity, however during home cage rest periods and intervals where different animals were in the two-photon box, animals were kept in the light condition (either light or darkness) consistent with their 12hr light/dark cycle.

### Two-photon image post-processing

Images were acquired using PrairieView acquisition software (Bruker) and converted into multi-page TIF files.

For spine imaging, registration and averaging was performed for each *z*-plane spanning the axial width of the dendrite to ensure all spines were captured across *z*-planes. A gaussian distribution was fit to the intensity of the dendrite, and the images were weighted according to the fit. Dendritic segments then underwent rigid global registration across days. Dendrite registration was performed using the MATLAB imregister function with regular-step gradient descent optimization and a mean square error metric configuration. The registered images underwent high-pass filtering to extract low amplitude spine features using code adapted from Suite2P’s enhanced mean image function^102^. The resulting ROIs were binarized using Otsu’s global threshold method for spine classification^103^. In most cases, the global threshold successfully isolated the single most prominent dendrite. In fields with higher background dendrites that were not desired, these extraneous dendrites were manually excluded. To identify spines that fall below the global threshold, the user manually specifies incrementally lower thresholds from which to select spines that were excluded in the initial binarization. Spines above the global threshold with an area of >1 μm^2^ were included in our analysis. To classify each spine as one of the four major morphological classes, we performed the following steps.

First, we found the base of the spine by identifying the region closest to the dendritic shaft. Second, we calculated the length of the spine by taking the Euclidean distance between the midpoint of the spine base and the most distant pixel. Third, this vector was divided evenly into three segments to find the spine head, neck, and base areas, respectively. Finally, spines were classified in the four categories, considering the following threshold parameters: stubby (neck length <0.2 μm and aspect ratio <1.3), thin (neck length >0.2 μm, spine length <0.7 μm, head circularity <0.8 μm), mushroom (neck length >0.2 μm, head circularity >0.8 μm), and filopodium (neck length >0.2 μm, spine length <0.8 μm, aspect ratio >1.3).

For calcium imaging sessions, the TIF files were processed using the Python implementation of Suite2P^102^. We briefly summarize their pipeline here. First, TIFs in the image stack undergo rigid registration using regularized phase correlations. This involves spatial whitening and then cross-correlating frames. Next, regions of interest (ROIs) are extracted by clustering correlated pixels, where a low-dimensional decomposition is used to reduce the size of the data. The number of ROIs is set automatically from a threshold set on the pixel correlations. We manually checked assigned ROIs based on location, morphology, and *DF*/*F* traces.

Since the hippocampal pyramidal cells are densely packed and the microperiscope reduces the axial resolution, we perform local neuropil subtraction using custom code (https://github.com/ucsb-goard-lab/two-photon-calcium-post-processing) to avoid neuropil contamination. The corrected fluorescence was estimated according to

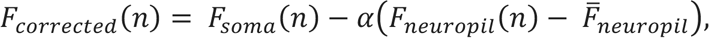

where *F_neuropil_* was defined as the fluorescence in the region <30 μm from the ROI border (excluding other ROIs) for frame *n*. *F_neuropil_* was *F_neuropil_* averaged over all frames. α was chosen from [0, 1] to minimize the Pearson’s correlation coefficient between *F_neuropil_* and *F_neuropil_* The Δ*F*/*F* for each neuron was then calculated as

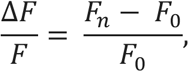

where *F*_*_ is the corrected fluorescence (*F_corrected_*) for frame *n* and *F*_/_ is defined as the first mode of the corrected fluorescence density distribution across the entire time series.

We deconvolved this neuropil subtracted Δ*F*/*F* to obtain an estimate for the instantaneous spike rate, which we used for the computation of neurons’ spatial information and mean event rate (fig. S4B, D). This inferred spike rate was obtained via a MATLAB implementation of a sparse, nonnegative deconvolution algorithm (OASIS) used for Ca^2+^ recordings^104^. We used an auto- regressive model of order 2 for the convolution kernel.

### Spine imaging data analysis

After rigid registration, high-pass filtering, and binarization of the dendritic segment, individual spines were extracted based on standard morphological criteria^105^. Spines projecting laterally from the dendritic segment were extracted and analyzed as individual objects, as described previously. The sum of the members of each spine class, as well as the total number of all spines, was recorded for each session. Spine totals (*S_total_*) were then broken down into 10 μm sections of the dendritic segment (*S_section_*) using the following calculation

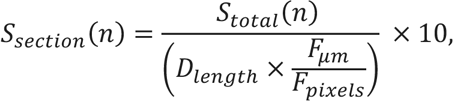

where length of the dendritic segment, *D_length_*, was determined by skeletonizing the dendritic shaft to 1 pixel in diameter, then taking the area of the pixels. *F_pixels_* is the FOV in pixels, which here was 760 pixels in each axis at 16× magnification, and *F_μm_*. is the FOV in µm, which was 52 μm in each axis.

Turnover was estimated at 12 hr increments; turnover here is defined as the net change in spines per session for each morphological class. Percent addition/subtraction, *T*, was calculated as

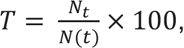

where *N*_t_ is spines that have been added or subtracted and *N*(*t*) is the total average number of spines. To determine the total population of spines on each dendrite, dendritic segments first underwent global registration across days, as described previously. After registration the centers of all spines across all days were overlaid, and spines falling under a spacing threshold of 1.7 μm were considered to be the same spine. The remaining ROIs represented the total cumulative population of dendritic spines across the time series.

Global registration was rigid so that spine shape would not be warped and misclassified as a different spine type, however small differences in dendritic morphology made the visualization of the exact same spine location difficult across days. Because of this, we also employed local registration, in which an 80 x 80 pixel ROI around the centroid of the dendritic spine was cut out and independently registered against the time series of images using the same registration parameters as in global registration. This is consistent with previous dendritic spine registration techniques^106^.

The coordinates of the locally registered spine ROI from the gaussian averaged image were applied to the same region on the binarized image, and a 40 x 40 pixel series of both average projection and binarized ROI images were displayed in a GUI format, along with recording number, so that the experimenter could manually confirm that the same spine was detected across days, and that the classification of the spine as either present or not present was correct (fig. S1). Manual curation of spines was performed blind to estrous stage.

Once spine numbers across recordings were confirmed and normalized to spines per 10 µm, recordings were interpolated to an archetypal cycle, where each stage was repeated twice and the cycle was repeated three times (i.e. D-D-P-P-E-E-M-M x3; Fig. 1F). Stages were first interpolated to a standard two-recording length, such that stages that lasted just one recording were repeated twice, and stages that lasted more than 2 recordings were averaged so that the first timepoint was the average of the first half of recordings within that stage (rounded up for stages with odd number recordings), and the second timepoint was the average of the second half. Stages lasting two recordings were left as-is. The resulting vector was input into the archetypal cycle such that the first observed stage determined the starting location of the vector. For instance, a recording beginning with proestrus would begin at the third slot in the archetypal cycle. This method was employed instead of circularizing the cycle to preserve the time course of turnover within dendrites.

To calculate the survival fraction curve *S*(*t*), we determined which spines were present at time *tn* that were not present at time *t0*^21, 47, 107^. For all spines this was considered any spine that was present on recording session 1 regardless of estrous stage. For proestrus-added spines, only spines for which the diestrus stage before proestrus as well as an entire cycle after proestrus were recorded were included in the analysis. Spines were considered to be proestrus-added when no spine was present in diestrus, regardless of when during the proestrus stage they appeared. The survival fraction of these spines was quantified such that recording 1 was the recording at which the spine appeared, and recording 2 was the first recording of the estrus stage immediately following proestrus. Survival fraction was quantified as

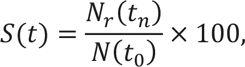

where *N_r_*(*t_n_*) are the total spines at *t_n_* that were also found in *t*_0_, and *N*(*t*_0_) are the total number of spines that were present in *t*_0_.

To calculate transition matrices (Fig. 2F), the population of spines present in diestrus, proestrus, and estrus were first classified, as well as the classifications of those spines in the stage immediately following (i.e. diestrus > proestrus, and proestrus > estrus). The transition matrices were calculated as the probability that a spine of a particular classification (e.g., mushroom) would transition to each of the other classifications (mushroom, stubby, thin, filipodia, no spine) during the stage transition. Matrices were pseudocolored to indicate transitions to more stable spine classes (red, above unity line), less stable spine classes (blue, below unit line) or no change (greyscale).

### Dendritic imaging data analysis

For dendritic imaging, mice were allowed to run head-fixed around the floating chamber for 15 minutes, during which they completed at least 10 but up to 20 laps. Visual and textural cues in the chamber were the same used for environment A in remapping analysis. The analysis of dendritic imaging data was broken down into 4 steps:

Find ROIs. Here, the user extracted ROIs using a custom GUI interface which overlaid the average projection on top of a pixel-wise activity map calculated using a modified kurtosis measure and asked the user to manually draw ROIs around the place cell of interest, which was highlighted in a separate window to guide ROI selection. Soma was selected using an elliptical ROI, and dendrite was selected using a freehand object. The somatic ROI was maintained for the rest of the analysis, but the dendritic ROI was automatically redrawn by dilating the skeletonized image defined by the outer bounds of the hand-drawn ROI with a structuring element 10-pixels in diameter. This helped to eliminate user error and standardize dendritic ROI selection. subROIs were created by splitting up the previously defined dendritic ROI into serial sections with a specified length, here 6 μm. A subset of somatic ROIs and their respective dendritic and sub-ROIs were identified as place cells according to the previously described pipeline, and put through the same next steps to determine estrous-dependent properties (fig. S2).

1. Extract ΔF/F. ΔF/F was calculated as is previously described in our place cell analysis, for soma ROIs, dendrite ROIs, branch ROIs, and subROIs.
2. Calculate co-tuning. Co-tuning between soma and dendrites and soma and subROIs were calculated using a standard Pearson’s correlation between ΔF/F traces extracted from their respective ROIs (Fig. 3G,L).
3. Calculate length constant. To determine length constant, bAPs were identified in somatic ΔF/F by extracting significantly elevated sections of the calcium trace^38^. Occasionally short sections of an AP fell below threshold, so gaps of < 5 frames (0.5 s) were interpolated to avoid counting single bAPs multiple times. During the AP event, subROI responses along the length of the dendrite were isolated and averaged, then normalized to the average somatic ΔF/F during the bAP interval. Each bAP was fit with an exponential decay function to determine the decay constant and goodness of fit. The exponential decay function was calculated as

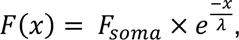

where *F_soma_* is the ΔF/F at the soma (normalized to 1), *x* is the distance in microns, and λ is the decay constant.

The cumulative population of bAPs had an averaged r^2^ of 0.81 ± 0.06 (mean ± sem), and a threshold r^2^ value of 0.7 was set to only consider good-quality bAPs. The length constant (λ), was calculated as the decay constant of the fit exponential decay function, and is equivalent to the distance a bAP travels before decaying to 37% of its initial amplitude. Length constants were capped at 400 µm, the maximum length of any dendrite in the dataset. The resulting bAP length constants were averaged across events within cells, and analyzed as a function of estrous stage (Fig. 3M).

### Population imaging data analysis

For calcium imaging experiments during exploration of the air-floated chamber, processed and synchronized behavioral data and 2P imaging data were used to identify place cells as follows.

First, the 1D track was divided into 72 equal bins (each ∼0.85 cm in length). Activity as a function of position (we refer to these as spatial tuning curves) was computed for each lap, with activity divided by occupancy of each binned location. We observed that in certain cases, the mice traversed the track at high speeds. To avoid misattribution of slow calcium signals to spatial bins (which were relatively small due to the length of the track), any lap where the average instantaneous speed was greater than 200 mm/s was removed and not considered for further analysis (an average of 6% of laps were removed). To assess the consistency of spatial coding of each cell, we randomly split the laps into two groups and computed the correlation coefficient between the averaged spatial tuning curves. We then did the same for shuffled data in which each lap’s spatial tuning curve was circularly permuted by a random number of bins.

Note that this was done for each lap, to avoid trivial effects that might emerge from circularly permuting data that was stereotyped along the track. This was performed 500 times, and the distribution of actual correlation coefficient values was compared to the distribution of circularly shuffled values using a two-sample Kolmogorov-Smirnov test (*α* = 0.01). The average correlation coefficient for actual data for each place cell was used as a metric to determine lapwise stability across estrous stages (fig. S4D). The distribution of these values had to pass a Cohen’s *D* analysis, having a score of greater than 1.2. A cell that passed these tests was considered a ‘consistent’ cell^42^.

To identify a neuron as a place cell, the neuron had to pass the consistency test, in addition to being fit well by a Gaussian function, 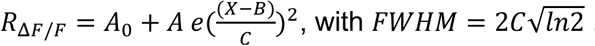. Note that in this convention, *C*^2^ = 2σ^2^. specifically, we required that: (1) the adjusted *R*^2^ > 0.375; (2) 2.5 cm < FWHM < 30.6 cm (50% of track length); 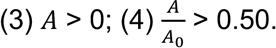. Cells that met these conditions were characterized as place cells; with place fields at the location of maximal activity and width defined as the FWHM. Note that these criteria are somewhat strict compared to traditional place field criteria. When tested with data in which individual laps were time shuffled, the approach yielded a false positive rate of 0%. In remapping experiments, only place cells that passed these qualifications in environment A were considered for further analyses.

To compute the spatial information of cell *j* (*SI*_*j*_; *112*), we used the following formula,

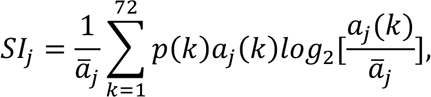

where α_*j*_ is the mean inferred spike rate of cell *j*, α_*j*_(*k*) is the mean inferred spike rate of cell *j* at position bin *k*, and *p*(*k*) is the probability of being at position bin *k*. We divide by α_*j*_ to have *SI*_*j*_ in units of bits/inferred spike (fig. S4B).

For single cell analysis, differences between peaks in the same place cell across three environments was measured in bins, then converted to circular distance in cm (Fig. 5C). For population vector analysis, population vectors were defined for each 0.85-cm bin of the lap-averaged tuning curves (72 bins in total) from all cells in the experimental group for each of the three environments. Pearson’s correlations were calculated between vectors of the same spatial bin across the three environments (A>B>A’) and averaged for every occurrence of the four stages within each animal. Because recordings in different stages had different numbers of cells, for recordings within each animal PVec correlations were bootstrapped across 100 iterations and n = 6 mice, sampling the minimum number of cells from each stage with replacement (Fig. 5D).

To further test estrous-dependent changes in spatial coding, we used a linear model to predict position in circular environments (B, A’) based on modeled firing rate distributions from a reference environment (A). For each iteration, cells were randomly sampled with replacement, using the minimum number of cells across stages for each animal (*n* = 577 neurons across 6 animals). We sampled 7 trials, the minimum number of trials across all recordings, from each environment for each cell. From the responses to environment A, we determined the firing rate distribution for each neuron. For each position in environments B and A’, the response of each neuron was multiplied by the Environment A firing rate distribution, and the scaled firing rate distribution were summed across all neurons and normalized to generate an estimated probability for each position. We then plotted the probability density of the estimated position at each actual position in environments B and A’, averaged over 100 iterations to account for sampling variability (Fig. 5E; fig. S6).

### Histology

Mice were euthanized with CO2 and transcardially perfused using 4% paraformaldehyde. Extracted brain tissue was immersion fixed overnight at 4°C. After 24 hrs, samples were moved to 1X PBS. Subsequently, 100 μm coronal sections were cut using a vibratome (Leica, Neuroscience Research Institute Microscopy Facility, UCSB). Sections were then mounted on gel subbed slides using Vectashield Antifade Mounting Medium with DAPI (Vector laboratories Inc; H-1200-10) and sealed under #0 coverslips. Images were taken using a Leica SP8 resonant scanning confocal at 64X magnification and grid stitched in ImageJ.

### Statistics

Spine turnover was evaluated in each time point using a nonparametric Wilcoxon rank sum test against a grand mean for each stage of the archetypal cycle (Fig. 1F; table S1), while in-text statistics were performed using a Wilcoxon rank sum test against a grand mean without interpolation (Fig. 1F). In experiments evaluating turnover as a function of spine type, linear mixed effects models were generated for each spine type, with fixed effects for stage and random effects for mouse (Fig. 1G; table S1). For dendritic activity comparisons, as well as comparisons between single place cell responses, linear mixed effects models were used with fixed effects for stage and random effects for mouse. Pairwise linear mixed effects models were used to evaluate contrasts between specific stages (n = 6 comparisons, Bonferroni corrected; Fig. 3G,L,M; Fig. 5C; fig. S2C,D; fig. S4; tables S2-6). Very small p-values (<10^−4^) were capped at *p* ≤ 10^−4^ as a lower bound on reasonable probabilities. In population comparisons where bootstrapping was employed, 95% confidence intervals were used to evaluate significance (Fig. 5D; fig. S6D). When performing bootstrapping, data were randomly sampled with replacement.

## Acknowledgements

We would like to thank Ikuko Smith, Catherine Wooley, and Karyn Frick for discussion of the results. We would also like to thank Claire Cady, Nathaniel Bailey, and Yifan Xu for assisting in cytology collection and staging, and Mounami Reddy Kayitha for assisting in data analysis.

## Authors contributions

Conceptualization: M.J.G., E.G.J., W.T.R, N.S.W.

Methodology: M.J.G., W.T.R, M.K., N.S.W.

Investigation: M.J.G., N.S.W.

Visualization: M.J.G., N.S.W.

Funding acquisition: M.J.G., N.S.W.

Project administration: M.J.G.

Supervision: M.J.G.

Writing – original draft: M.J.G., N.S.W.

Writing – review & editing: M.J.G., E.G.J., W.T.R, N.S.W.

## Data and materials availability

Spine morphology images and processed calcium imaging traces generated in this study have been deposited in a public repository (URL to be added upon acceptance). The code used to measure spine morphology, dendritic activity, and place cell dynamics are available at (URL to be added upon acceptance).

## Supplementary Materials

Figs. S1 to S6 Tables S1 to S6 Movies S1 to S3

---

**Supplementary Figure 1.**
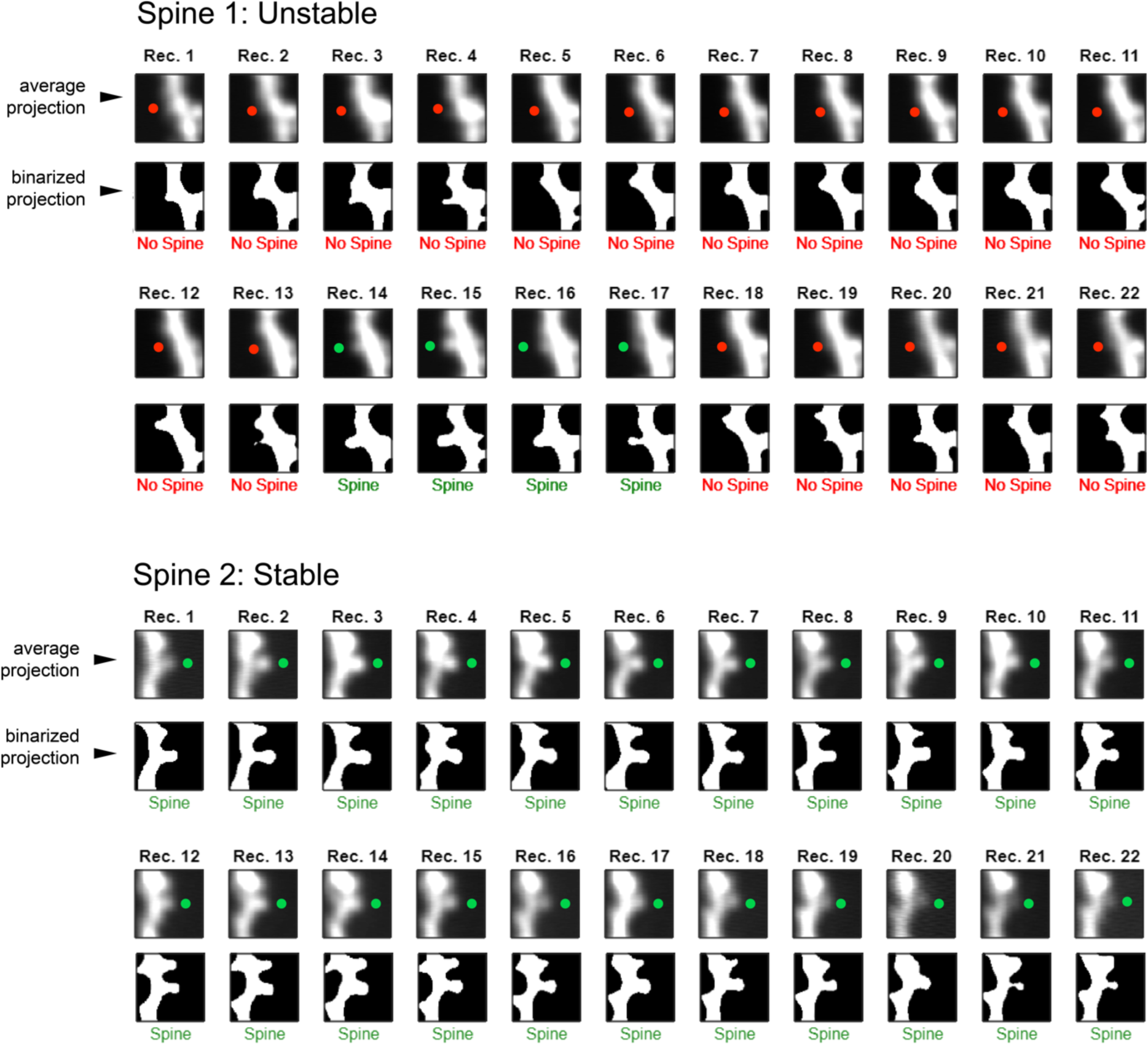
The spine detection GUI catches outlier false classifications. Here two examples of the spine detection GUI are shown, one where a spine was initially pruned then grew back after several recordings (unstable), and one where the spine was present across all recordings (stable). For each case, the first row of images is a gaussian-averaged projection, while the second row is a binarized and thresholded projection. Colored circles indicate the location and classification for each spine (green = spine, red = no spine). Here all classifications are correct, however for outlier cases in which a classification is incorrect (i.e. spine when there was no spine, or no spine when there was a spine), the user has the option to manually adjust the classification. The output data structure records where changes were made.

**Supplementary Figure 2.**
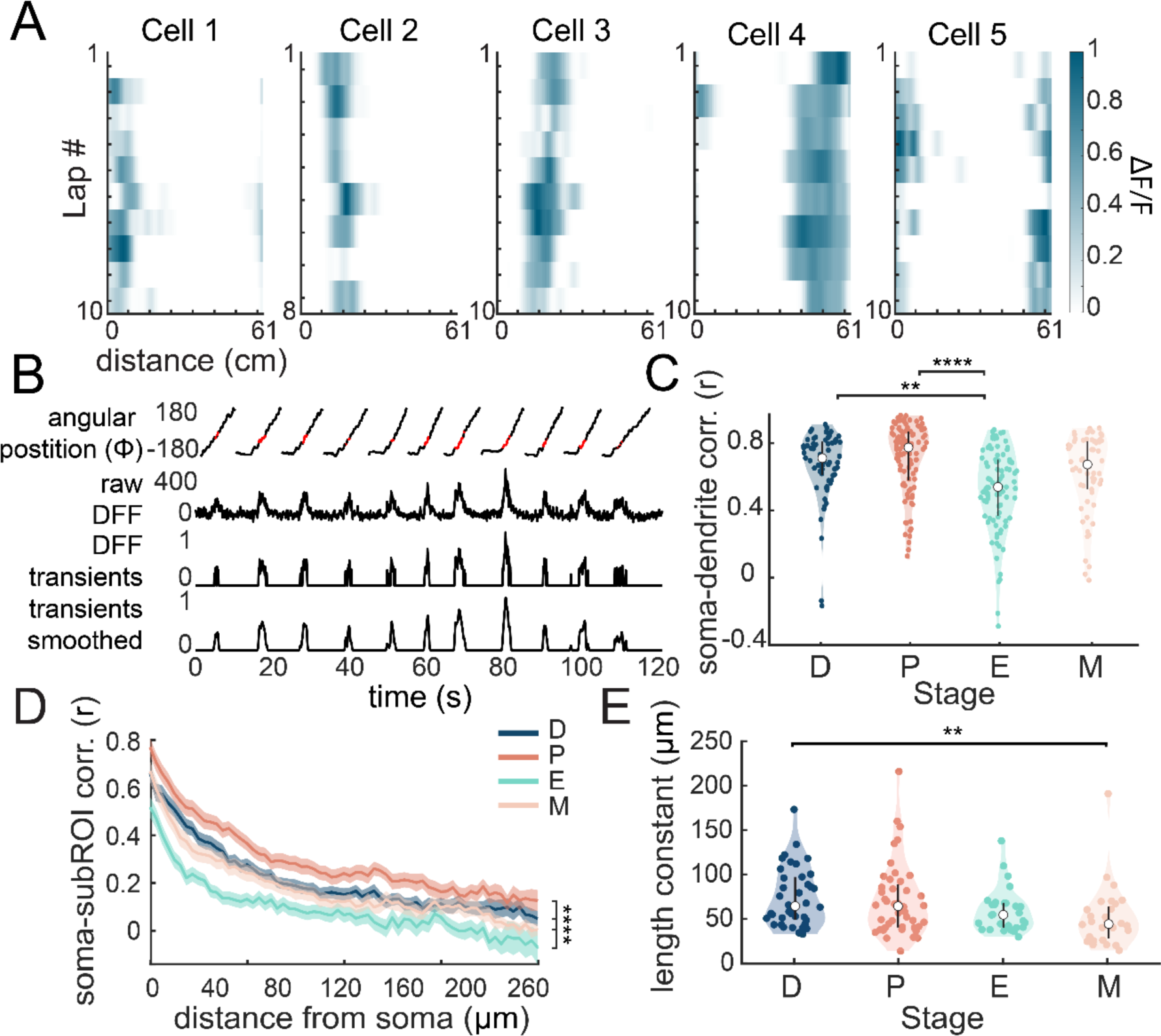
Estrous modulates the somatodendritic and intradendritic activity of place cells while a mouse is in motion. A. Calcium processing pipeline for an example place cell. Raw ΔF/F is thresholded and rectified and the ΔF/F transient trace is smoothed to remove noise. Regions of the angular position output of the Neurotar that correspond to when the cell is firing are labeled in red. B. Example place cell smoothed ΔF/F transients across laps through the floating chamber. C. Correlations between somatic and dendritic place cell ΔF/F from the entire recording across estrous stages. Mean ± standard error. Pairwise linear mixed effect model. \*\**p* < 0.01, \*\*\*\**p* < 0.0001. D. Correlations between place cell somas and their increasingly distal subROIs along the dendritic branch across estrous stages. Mean ± standard error. Bar indicates significant modulation across all stages, linear mixed effects model. \*\*\*\**p* < 0.0001. E. The distribution of length constants from bAPs for every place cell recorded across the estrous cycle. Mean ± standard error. Pairwise linear mixed effect model. \*\**p* < 0.01.

**Supplementary Figure 3.**
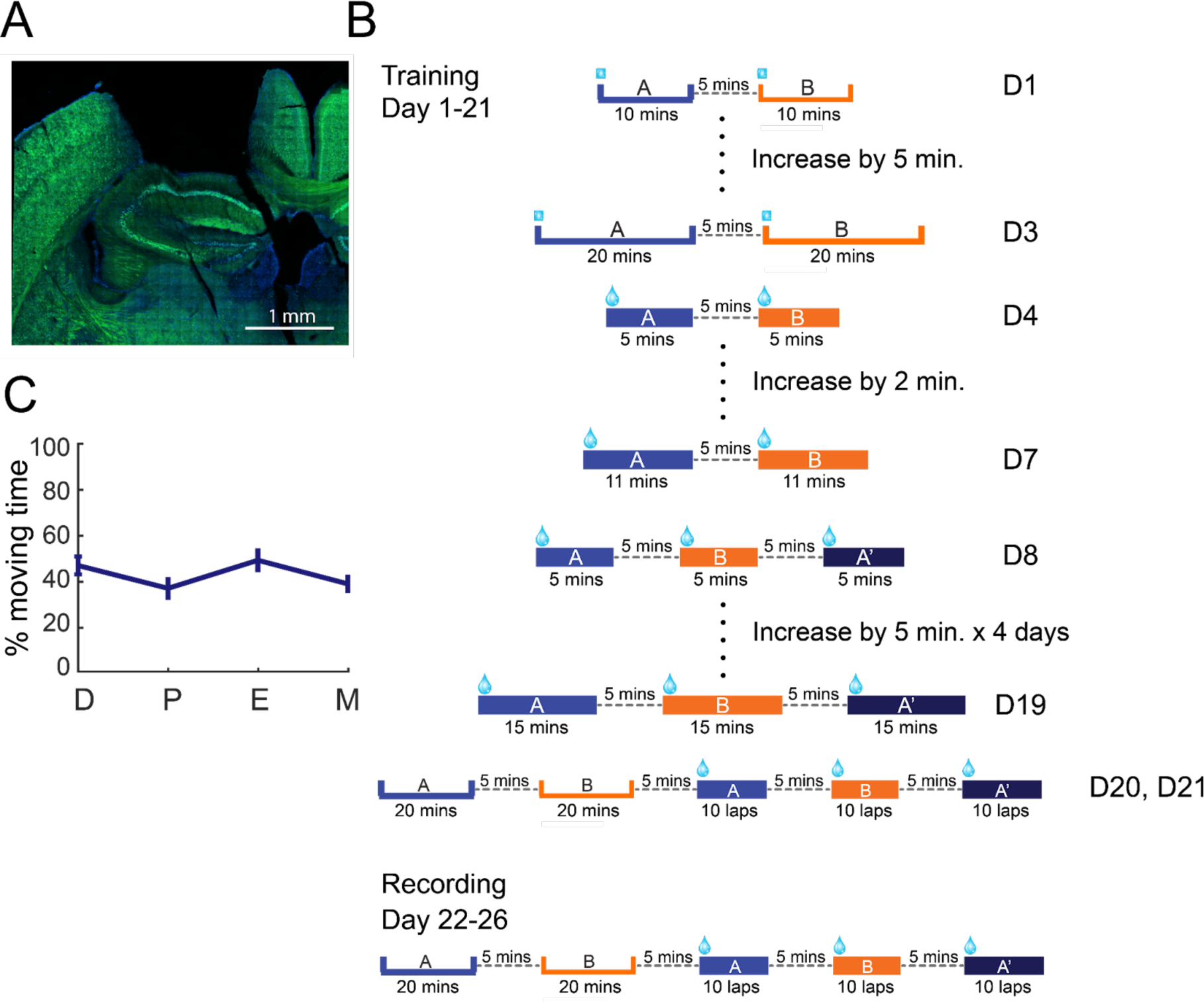
Behavioral training schematic. A. Confocal image of a CaMKII-Cre x TIT2L-GCaMP6s coronal section stained with DAPI illustrating the location of the glass plug above CA1. B. Training and recording schematic for remapping experiments. Training occurred over the course of 21 days with gradual acclimation to the full three rounds of head fixation. Open boxes indicate a freely moving mouse, filled boxes indicate a head fixed mouse. C. Percent moving time across stages during the recording phase. After several weeks of behavioral task training, percent moving time reaches a steady state across stages. Mean ± standard error.

**Supplementary Figure 4.**
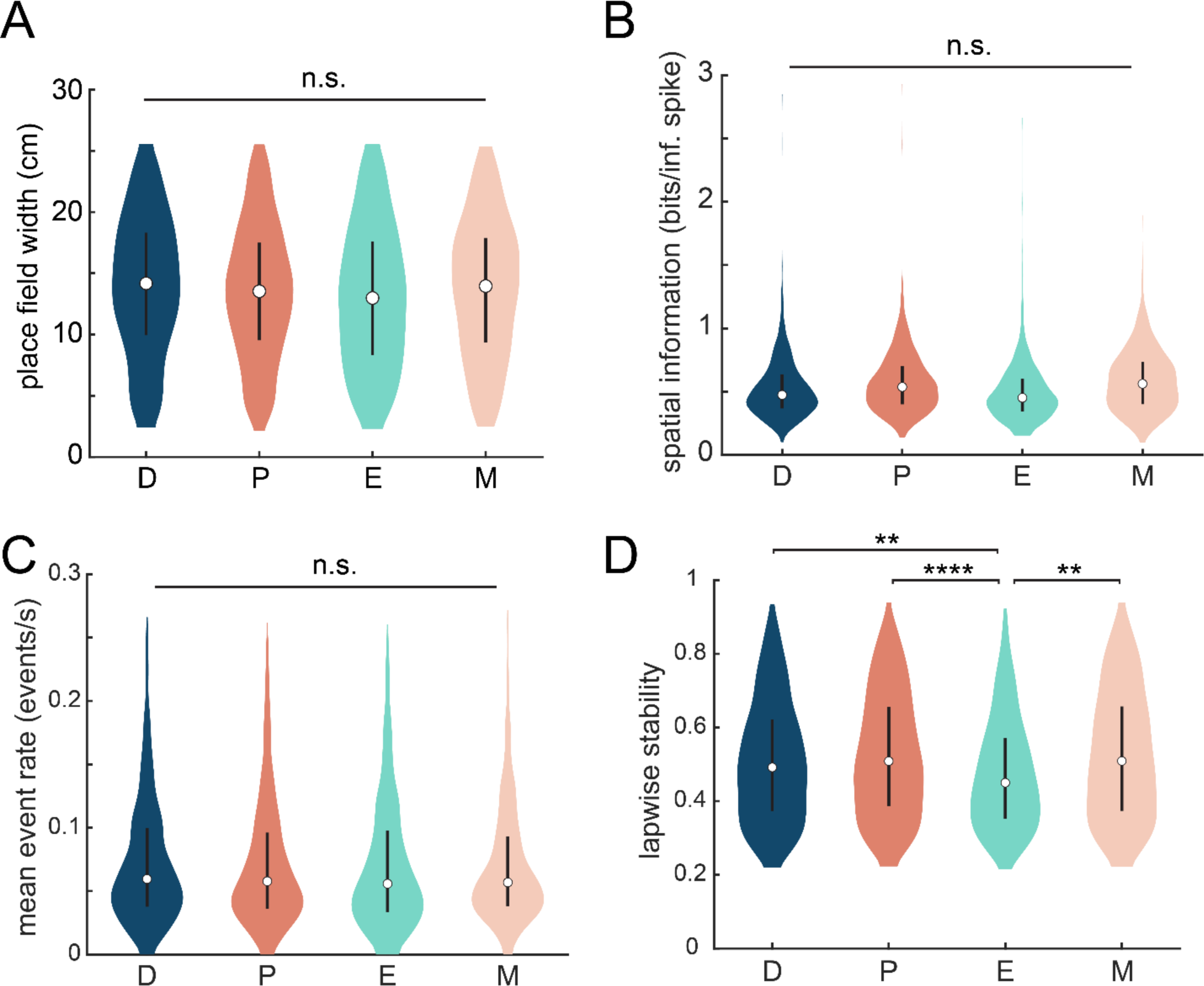
Place cell firing characteristics are largely stable across the estrous cycle. A. Place field width (FWHM) distributions across stages. Mean ± standard error. Linear mixed-effects model, n.s. = not significant. B. Spatial information score in bits/inferred spike across stages. Mean ± standard error. Linear mixed-effects model, n.s. = not significant. C. Mean event rate across stages (events/s). Mean ± standard error. Linear mixed-effects model, n.s. = not significant. D. Lapwise stability, calculated as correlations between two halves of lap data across stages. Pairwise linear mixed-effects model. \*\**p* < 0.01, \*\*\*\**p* < 0.0001.

**Supplementary Figure 5.**
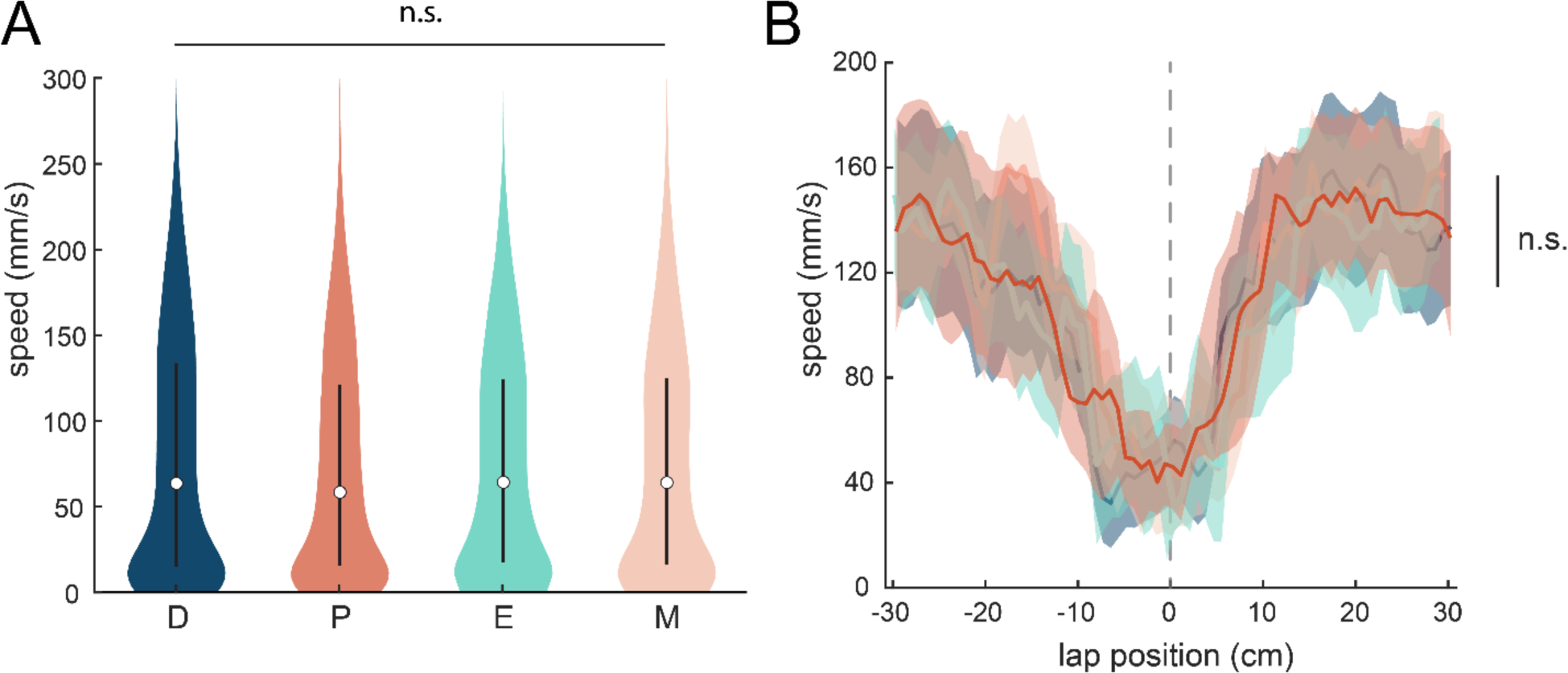
Behavioral parameters are not significantly modulated by estrous cycle stage. A. Distributions of average speed for each recording across n=6 animals as a function of estrous stage. Mean ± standard error. Pairwise linear mixed-effects models. n.s. = not significant. B. Anticipatory slowing behavior before and after reward administration (vertical line) for *n* = 56 recordings and n = 6 mice. Mean ± standard error. Pairwise linear mixed-effects models. n.s. = not significant.

**Supplementary Figure 6.**
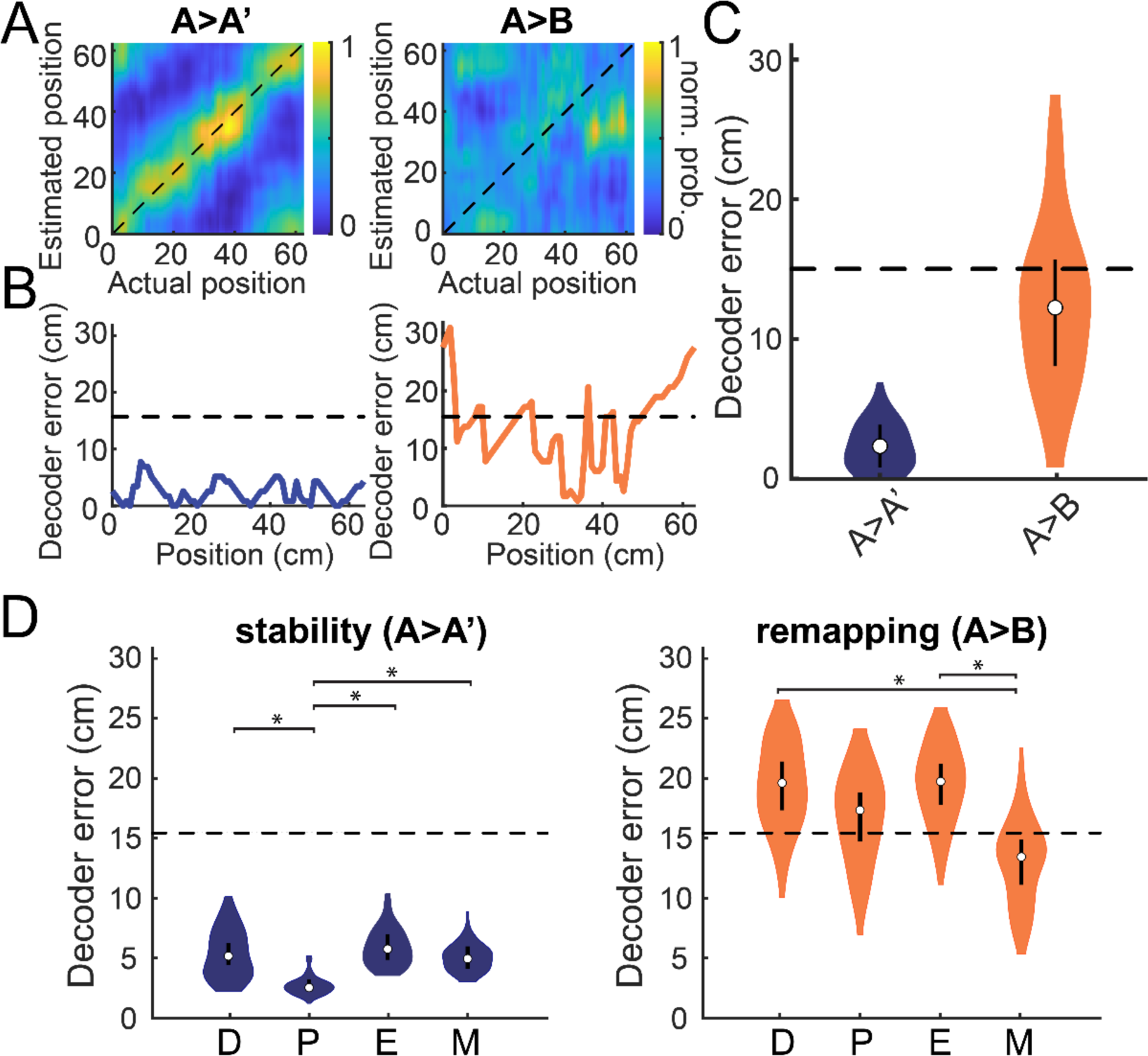
Population-level spatial coding is modulated by the estrous cycle. A. Probability density plot of a linear decoder trained on the place cell responses of environment A and used to estimate position in environment A’ (left) and environment B (right). Accurate estimates will fall along the unity line (dashed black diagonal). B. Prediction error in cm along the circular track for a decoder trained on environment A predicting position in environment A’ (purple) compared to environment B (orange). C. The distribution of prediction error between the same environments (A>A’, blue) compared to different environments (A>B, orange) in a single recording. Mean ± standard error. D. The distribution of prediction error of the decoder tested across estrous stages (D = diestrus, P = proestrus, E = estrus, M = metestrus), between the same environments (A>A’, stability) compared to different environments (A>B; remapping). Distributions were created by bootstrapping data from n = 6 mice over 100 iterations, sampling with replacement. Mean ± bootstrapped 95% CI. Asterisks denote significance. Significance was defined as lower 5% CI higher than higher 95% CI.

**Supplementary Table 1.** Complete statistics for Fig. 1F (Wilcoxon signed-rank test against a grand mean for all points) and Fig. 1G (pairwise linear mixed effect models Bonferroni-corrected for multiple comparisons). Lower bound capped at 10^-4^, upper bound capped at 1.0. α = 0.05.

| Comparison | *p* value | Test Statistic |
| --- | --- | --- |
| Figure 1F |  | W |
| P1 | 0.0840 | 45 |
| P2 | 0.0039 | 54 |
| E1 | 0.0002 | 0 |
| E2 | 0.0002 | 0 |
| M1 | 0.9405 | 107 |
| M2 | 0.6542 | 117 |
| D1 | 0.7652 | 113 |
| D2 | 0.2790 | 76 |
| P3 | 0.0001 | 230 |
| P4 | 0.0001 | 231 |
| E3 | 0.0001 | 5 |
| E4 | 0.0001 | 4 |
| M3 | 0.9405 | 107 |
| M4 | 0.7652 | 97 |
| D3 | 0.7089 | 95 |
| D4 | 0.8519 | 100 |
| P5 | 0.0001 | 105 |
| P6 | 0.0001 | 105 |
| E5 | 0.0625 | 0 |
| E6 | 0.0625 | 0 |

**Supplementary Table 1.** Complete statistics for Fig. 1F (Wilcoxon signed-rank test against a grand mean for all points) and Fig. 1G (pairwise linear mixed effect models Bonferroni-corrected for multiple comparisons). Lower bound capped at $10^{-4}$ , upper bound capped at 1.0. $\alpha = 0.05$ .
|  |  |  |
| --- | --- | --- |
| <u>Filopodium overall</u> | 0.1948 | 1.60(3) |
| Filopodium (D-P) | 1.0000 | 0.71(1) |
| Filopodium (D-E) | 1.0000 | 0.66(1) |
| Filopodium (D-M) | 1.0000 | 0.96(1) |
| Filopodium (P-E) | 0.4928 | 3.18(1) |
| Filopodium (P-M) | 0.2924 | 4.13(1) |
| Filopodium (E-M) | 1.0000 | 0.02(1) |
| <u>Thin overall</u> | 0.0019 | 5.41(3) |
| Thin (D-P) | 0.0676 | 7.06(1) |
| Thin (D-E) | 1.0000 | 1.41(1) |
| Thin (D-M) | 1.0000 | 0.00(1) |
| Thin (P-E) | 0.0007 | 18.41(1) |
| Thin (P-M) | 0.1144 | 5.97(1) |
| Thin (E-M) | 1.0000 | 1.28(1) |
| <u>Stubby overall</u> | 0.0497 | 2.72(3) |
| Stubby (D-P) | 0.7190 | 2.53(1) |
| Stubby (D-E) | 1.0000 | 1.79(1) |
| Stubby (D-M) | 1.0000 | 0.05(1) |
| Stubby (P-E) | 0.1030 | 6.18(1) |
| Stubby (P-M) | 1.0000 | 1.29(1) |
| Stubby (E-M) | 1.0000 | 1.73(1) |
| <u>Mushroom overall</u> | 0.8232 | 0.30(3) |
| Mushroom (D-P) | 1.0000 | 0.15(1) |
| Mushroom (D-E) | 1.0000 | 0.82(1) |
| Mushroom (D-M) | 1.0000 | 0.25(1) |
| Mushroom (P-E) | 1.0000 | 0.29(1) |
| Mushroom (P-M) | 1.0000 | 0.02(1) |
| Mushroom (E-M) | 1.0000 | 0.17(1) |

**Supplementary Table 2.**
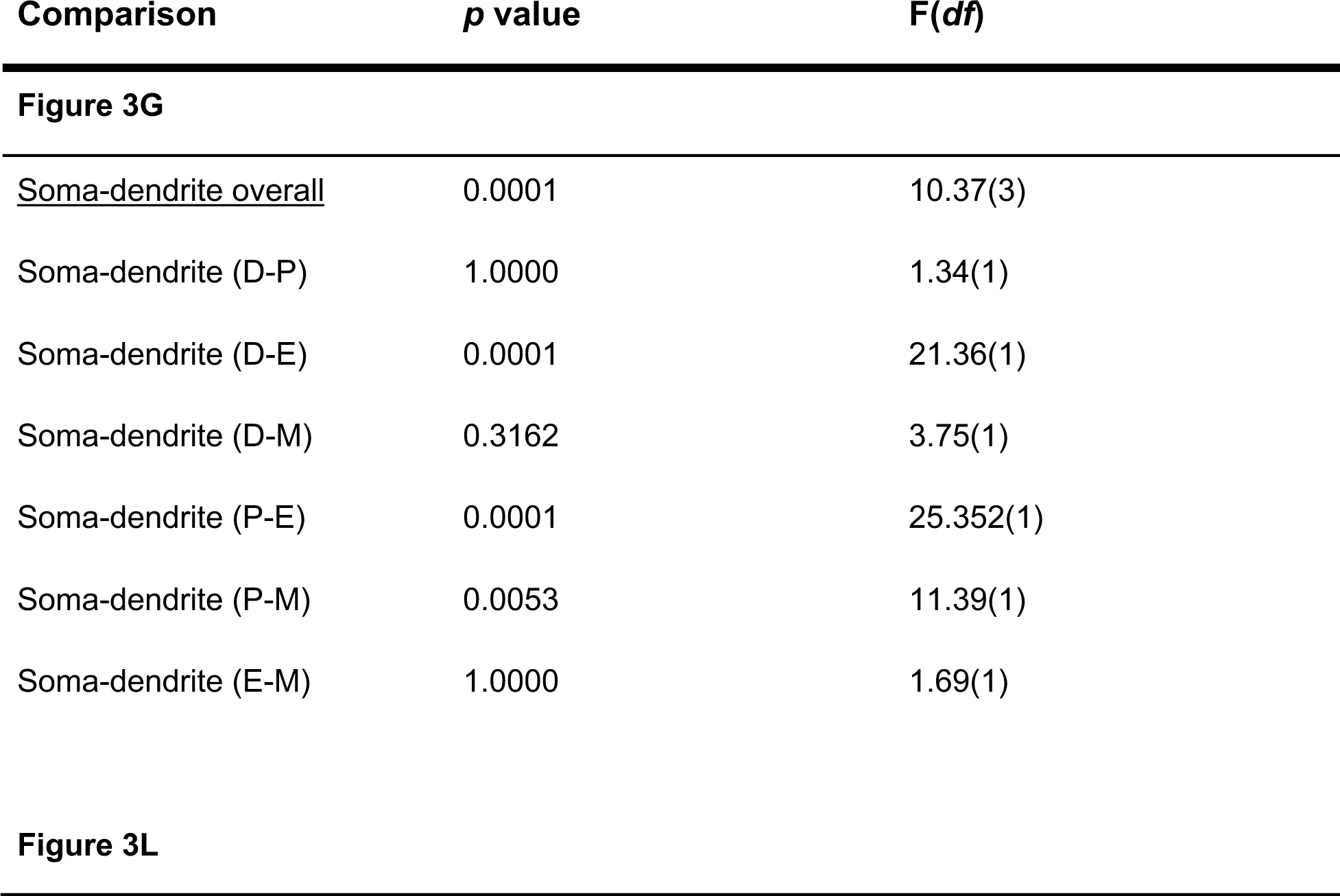
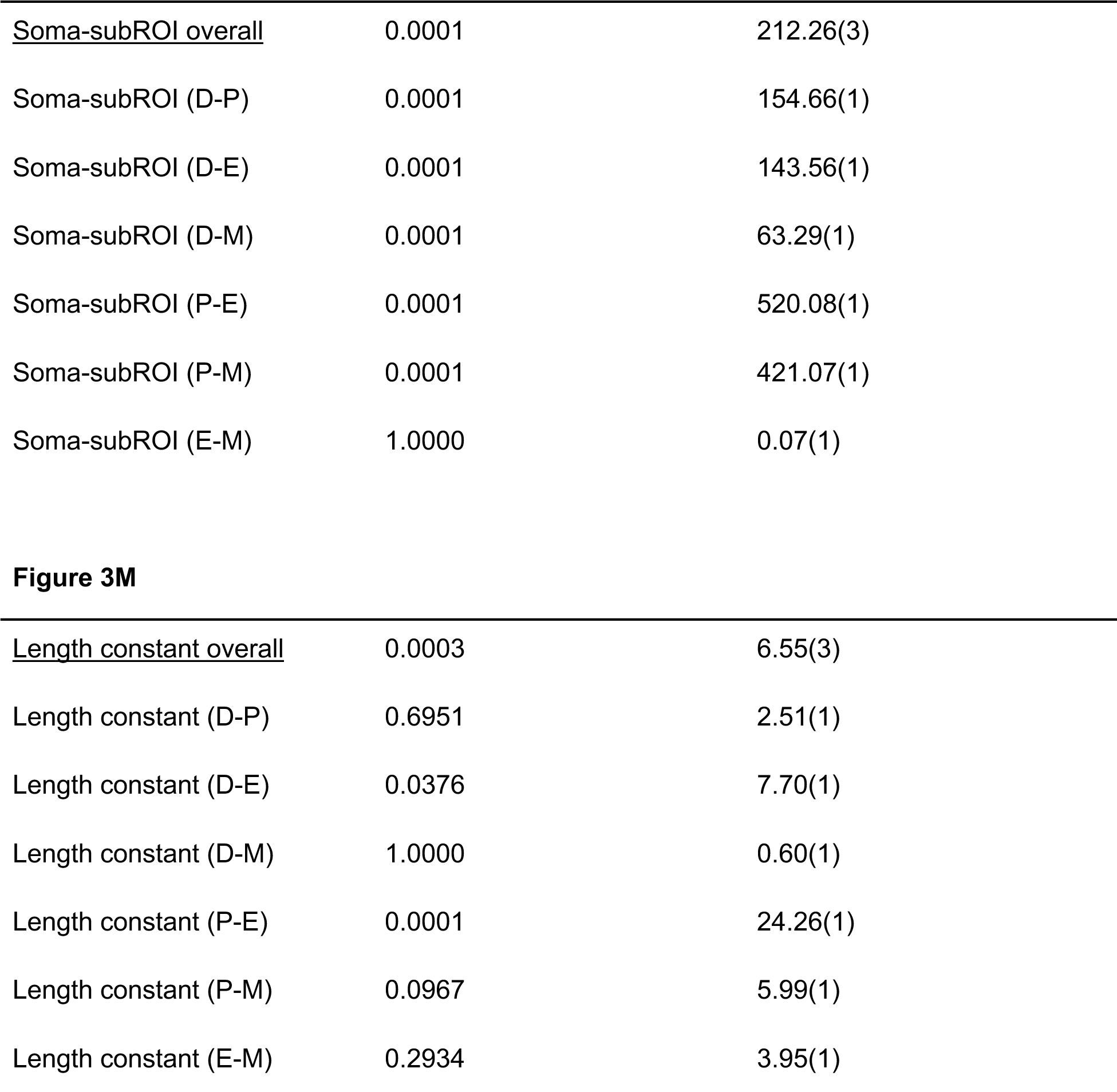
Complete statistics for Fig. 3G, L, and M. Pairwise linear mixed effect models Bonferroni-corrected for multiple comparisons. Lower bound capped at 10^-4^, upper bound capped at 1.0. α = 0.05.

**Supplementary Table 3.**
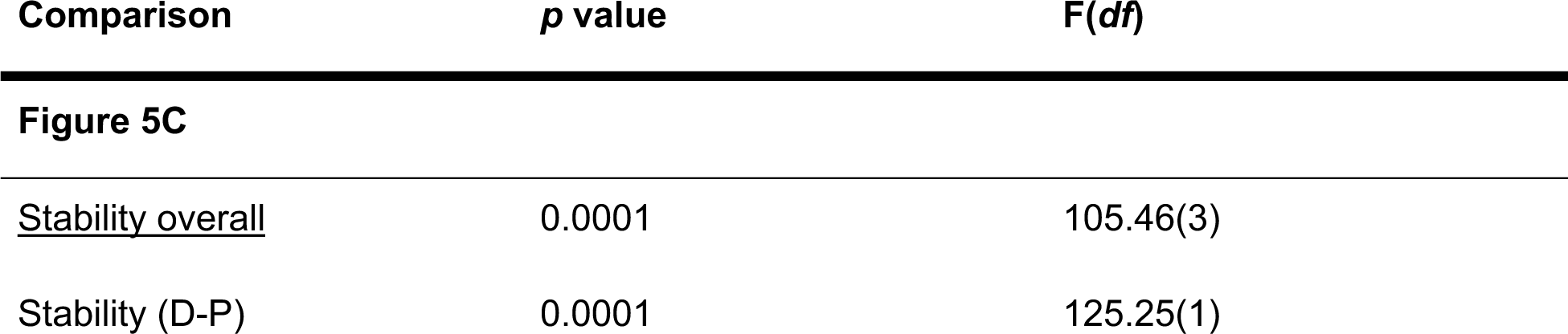
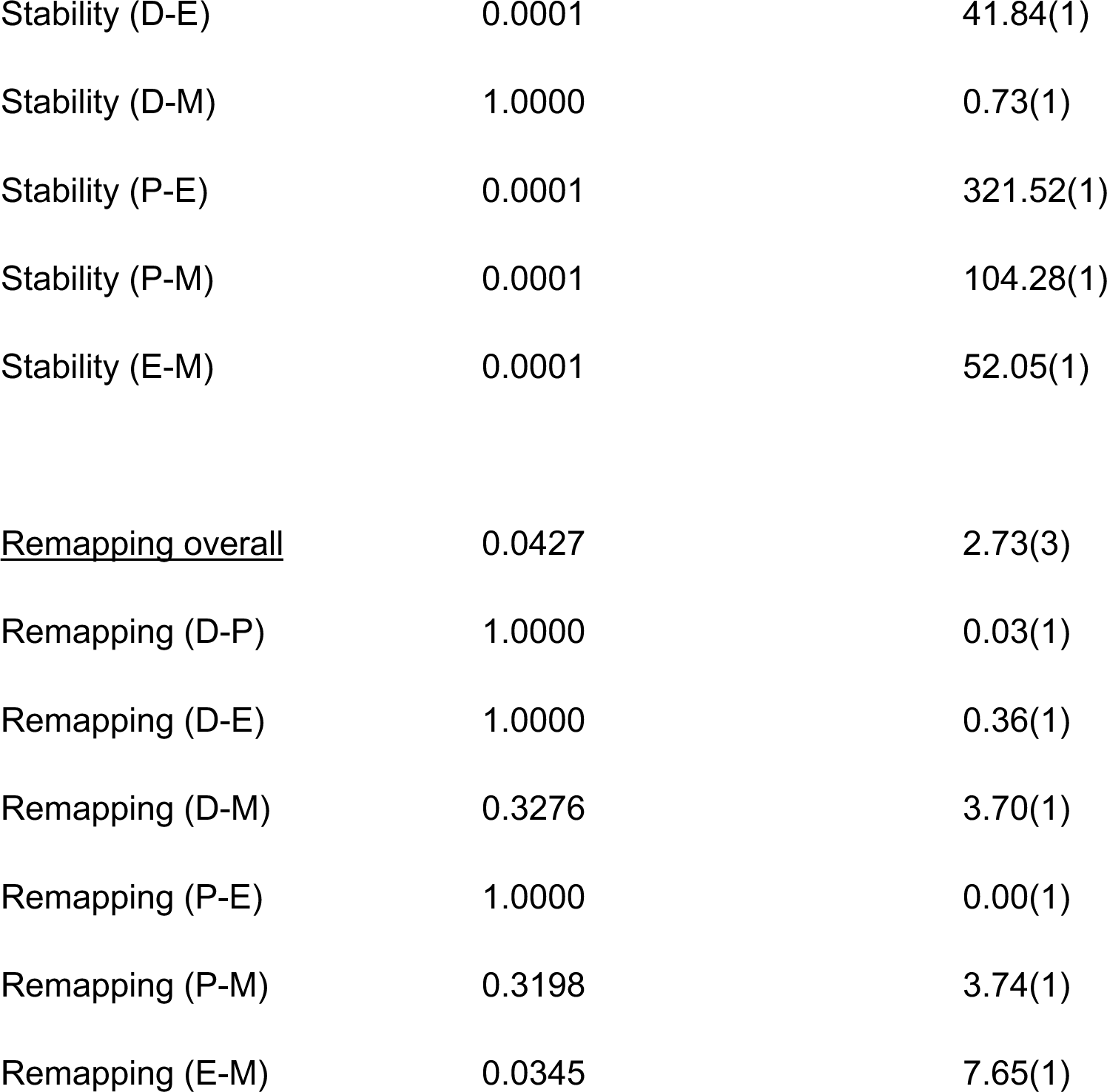
Complete statistics for Fig. 5C. Pairwise linear mixed effect models Bonferroni-corrected for multiple comparisons. Lower bound capped at 10^-4^, upper bound capped at 1.0. α = 0.05.

**Supplementary Table 4.**
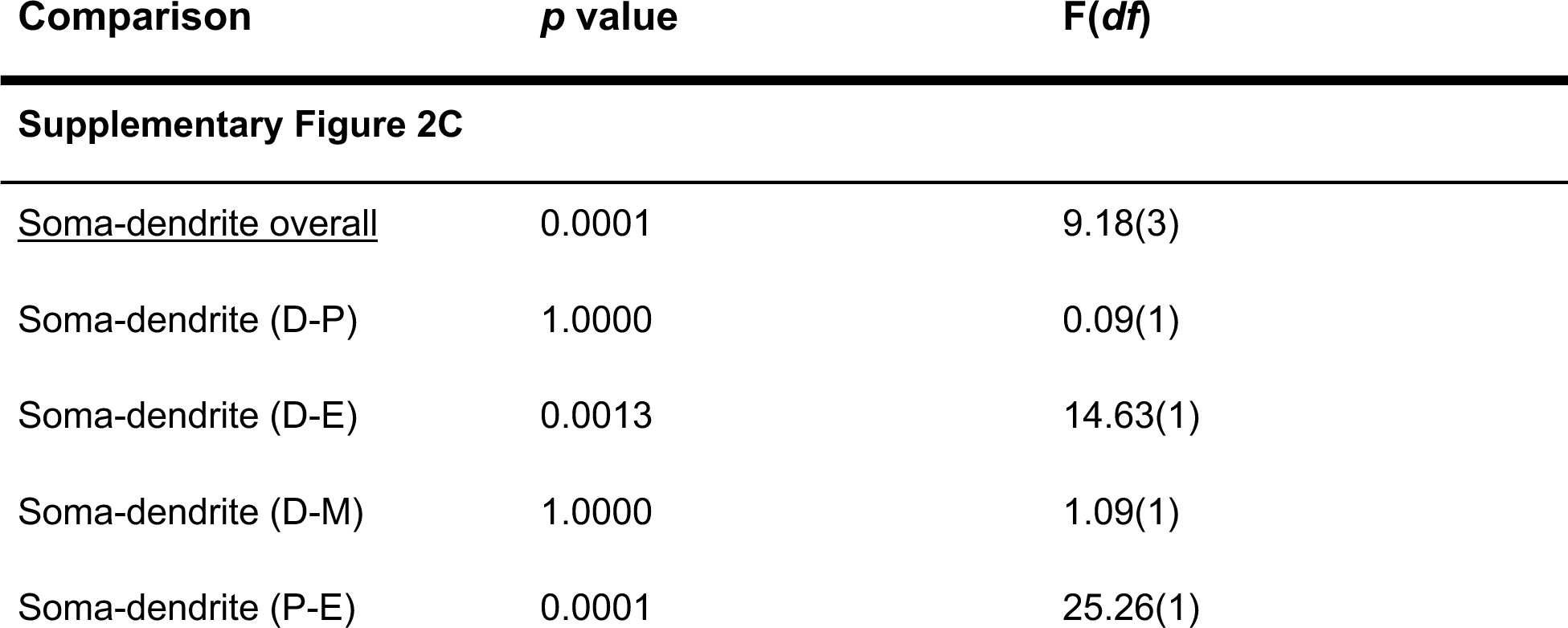
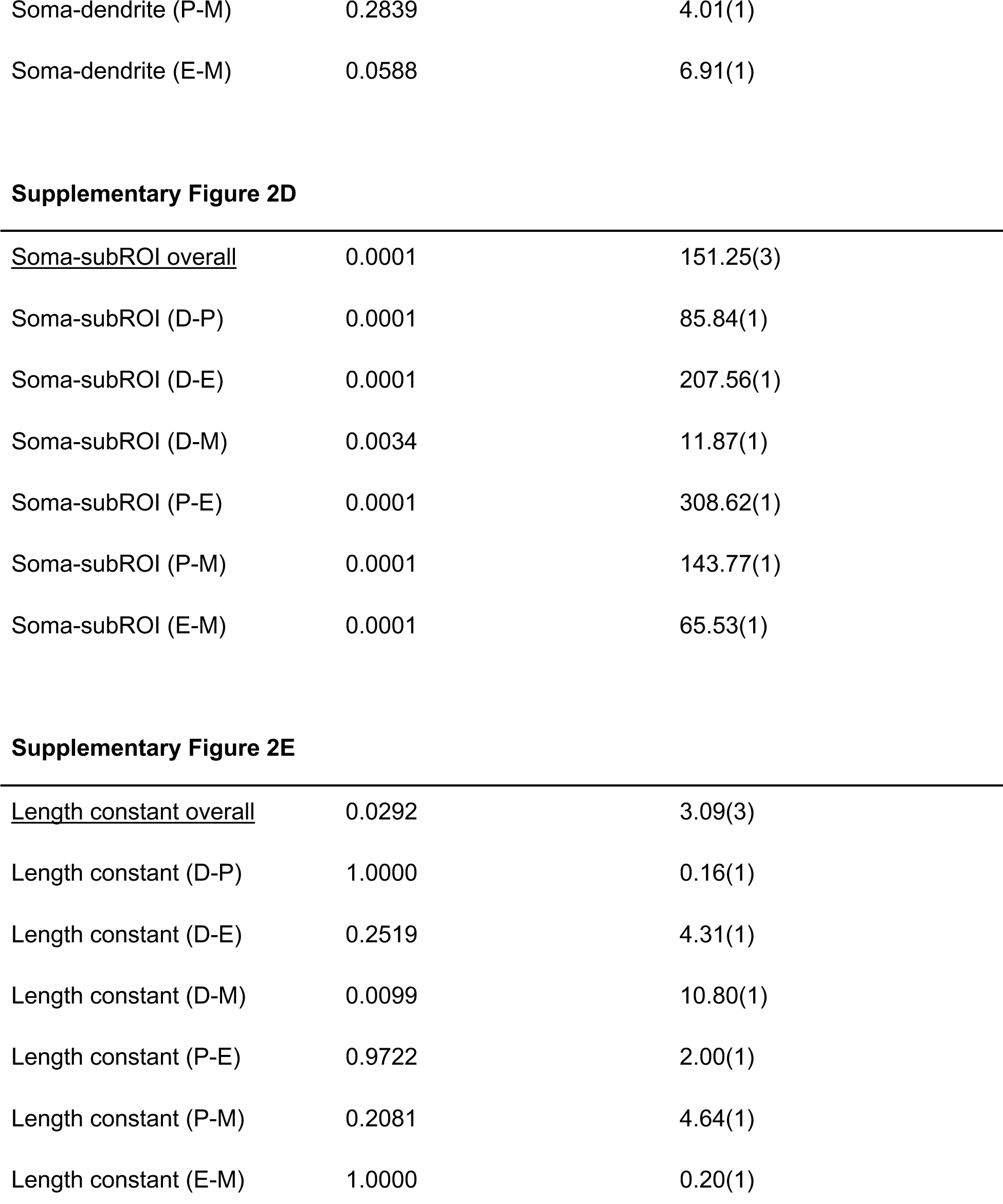
Complete statistics for Supplementary Figure 2C-E. Pairwise linear mixed effect models Bonferroni-corrected for multiple comparisons. Lower bound capped at 10^-4^, upper bound capped at 1.0. α = 0.05.

**Supplementary Table 5.** Complete statistics for Supplementary Figure 4. Pairwise linear mixed effect models Bonferroni-corrected for multiple comparisons. Lower bound capped at 10^-4^, upper bound capped at 1.0. α = 0.05.

| Comparison | *p* value | F(*df*) |
| --- | --- | --- |
| **Supplementary Figure 4A** |  |  |
| <u>Place field width overall</u> | 0.1375 | 1.36(3) |
| Place field width (D-P) | 1.0000 | 0.27(1) |
| Place field width (D-E) | 0.1047 | 1.35(1) |
| Place field width (D-M) | 1.0000 | 0.25(1) |
| Place field width (P-E) | 1.0000 | 0.48(1) |
| Place field width (P-M) | 1.0000 | 0.01(1) |
| Place field width (E-M) | 1.0000 | 0.33(1) |
| **Supplementary Figure 4B** |  |  |
| <u>Spatial information overall</u> | 0.2457 | 0.92(3) |
| Spatial information (D-P) | 0.1629 | 2.66(1) |
| Spatial information (D-E) | 0.0972 | 3.67(1) |
| Spatial information (D-M) | 0.5196 | 1.70(1) |
| Spatial information (P-E) | 0.0614 | 4.55(1) |
| Spatial information (P-M) | 1.0000 | 0.08(1) |
| Spatial information (E-M) | 0.0745 | 3.82(1) |
| **Supplementary Figure 4C** |  |  |
| <u>Event rate overall</u> | 0.1196 | 1.13(3) |
| Event rate (D-P) | 0.0754 | 3.23(1) |
| Event rate (D-E) | 0.0545 | 4.14(1) |
| Event rate (D-M) | 1.0000 | 0.67(1) |
| Event rate (P-E) | 0.0604 | 3.80(1) |
| Event rate (P-M) | 0.0678 | 3.89(1) |
| Event rate (E-M) | 0.5458 | 1.04(1) |

**Supplementary Table 5.** Complete statistics for Supplementary Figure 4. Pairwise linear mixed effect models Bonferroni-corrected for multiple comparisons. Lower bound capped at $10^{-4}$ , upper bound capped at 1.0. $\alpha = 0.05$ .
|  |  |  |
| --- | --- | --- |
| <u>Stability overall</u> | 0.0001 | 17.06(3) |
| Stability (D-P) | 0.0605 | 5.88(1) |
| Stability (D-E) | 0.0029 | 12.22(1) |
| Stability (D-M) | 0.0547 | 7.63(1) |
| Stability (P-E) | 0.0001 | 54.62(1) |
| Stability (P-M) | 1.0000 | 0.55(1) |
| Stability (E-M) | 0.0014 | 13.58(1) |

**Supplementary Table 6.**
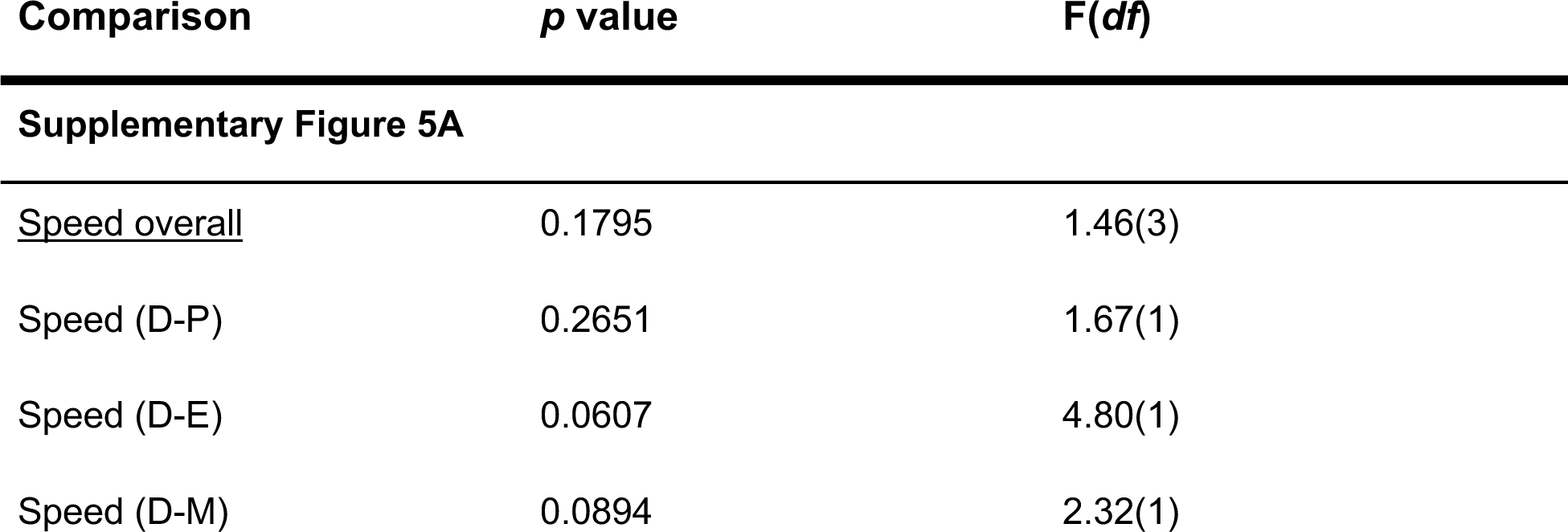
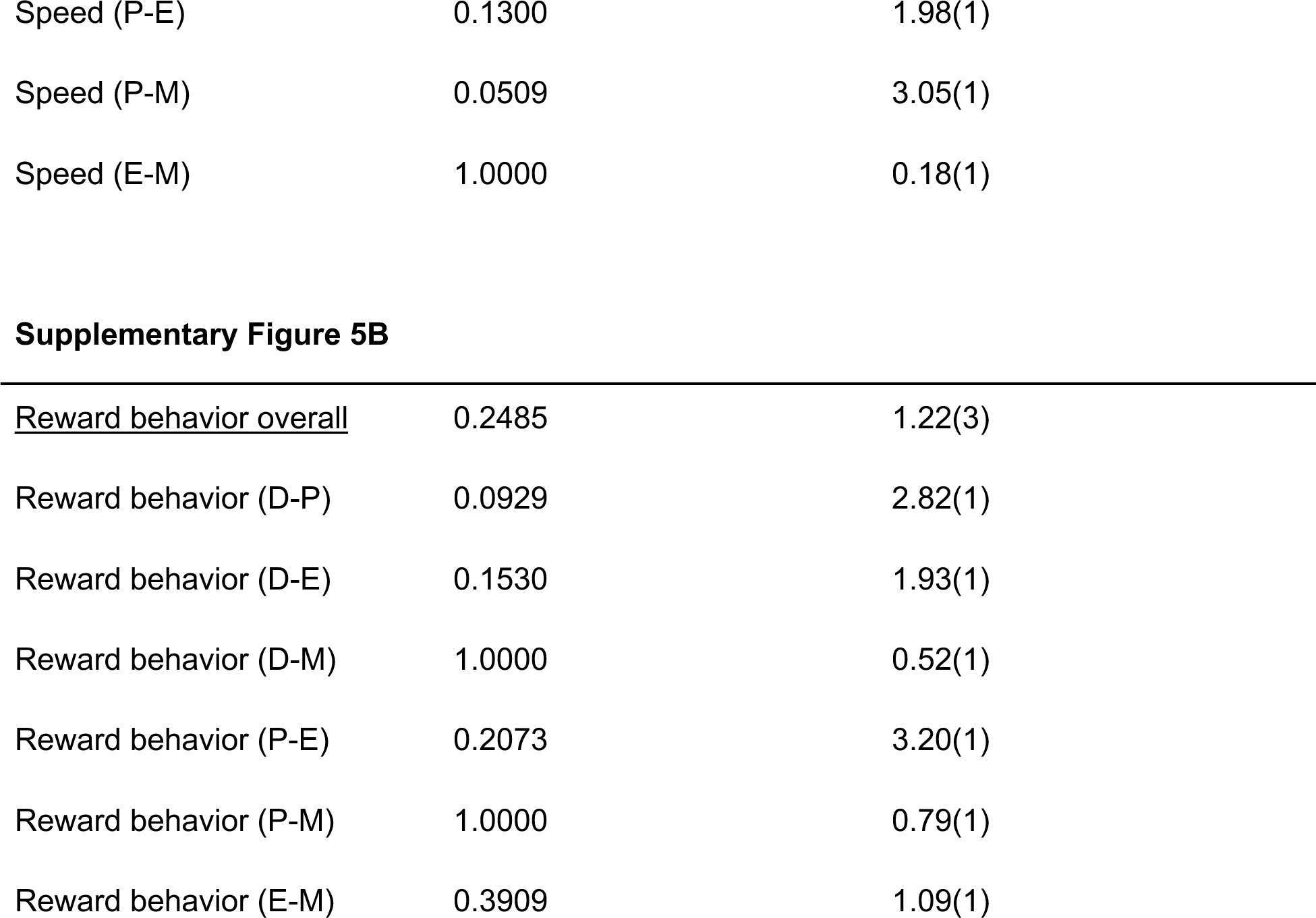
Complete statistics for Supplementary Figure 5. Pairwise linear mixed effect models Bonferroni-corrected for multiple comparisons. Lower bound capped at 10^-4^, upper bound capped at 1.0. α = 0.05.

| Comparison | *p* value | F(df) |
| --- | --- | --- |
| Supplementary Figure 5A |  |  |
| <u>Speed overall</u> | 0.1795 | 1.46(3) |
| Speed (D-P) | 0.2651 | 1.67(1) |
| Speed (D-E) | 0.0607 | 4.80(1) |
| Speed (D-M) | 0.0894 | 2.32(1) |
| Speed (P-E) | 0.1300 | 1.98(1) |
| Speed (P-M) | 0.0509 | 3.05(1) |
| Speed (E-M) | 1.0000 | 0.18(1) |

**Supplementary Table 6.** Complete statistics for Supplementary Figure 5. Pairwise linear mixed effect models Bonferroni-corrected for multiple comparisons. Lower bound capped at $10^{-4}$ , upper bound capped at 1.0. $\alpha = 0.05$ .
|  |  |  |
| --- | --- | --- |
| <u>Reward behavior overall</u> | 0.2485 | 1.22(3) |
| Reward behavior (D-P) | 0.0929 | 2.82(1) |
| Reward behavior (D-E) | 0.1530 | 1.93(1) |
| Reward behavior (D-M) | 1.0000 | 0.52(1) |
| Reward behavior (P-E) | 0.2073 | 3.20(1) |
| Reward behavior (P-M) | 1.0000 | 0.79(1) |
| Reward behavior (E-M) | 0.3909 | 1.09(1) |

**Supplementary Movie 1.** Recording of somatodendritic activity through the microperiscope implant.

**Supplementary Movie 2.** Mouse navigating the air-floated chamber while head-fixed.

**Supplementary Movie 3.** Recording of CA1 through plug implant.

